# Egocentric value maps of the near-body environment

**DOI:** 10.1101/2022.08.18.504456

**Authors:** Rory John Bufacchi, Richard Somervail, Aoife Maria Fitzpatrick, Roberto Caminiti, Gian Domenico Iannetti

## Abstract

Body-part centric response fields are pervasive: they are observed in single neurons, fMRI, EEG, and multiple behavioural measures. This prevalence across scales and measures makes them excellent candidates for studying systems-level neuroscience. Nonetheless, they remain poorly understood because we lack a unifying formal explanation of their origins and role in wider brain function. Here, we provide such explanation.

We use reinforcement learning to analytically explain the existence of body-part centric receptive fields, also known as peripersonal field. We then simulate multiple experimental findings considered foundational in the peripersonal space literature. Our results demonstrate that peripersonal fields naturally arise from two simple and plausible assumptions about living agents: 1) they experience reward when they contact objects in the environment, and 2) they act to maximise reward. These simple assumptions are enough to explain empirical findings on stimulus kinematics, tool use, valence, and network-architecture.

Our explanation provides further insight. First, it offers multiple empirically testable predictions. Second, it offers a formal description of the notion that the world-agent state is encoded in parieto-premotor cortices, using motor primitives: peripersonal fields provide building blocks that together create a short-term model of the world near the agent in terms of its future states; a successor representation. This short-term, close-range egocentric peripersonal map is analogous to the long-term, long-range allocentric spatial map of place and grid cells, which underlie locomotion and navigation to reach distant objects. Together, these allocentric and egocentric maps allow efficient interactions with a changing environment across multiple spatial and temporal scales.

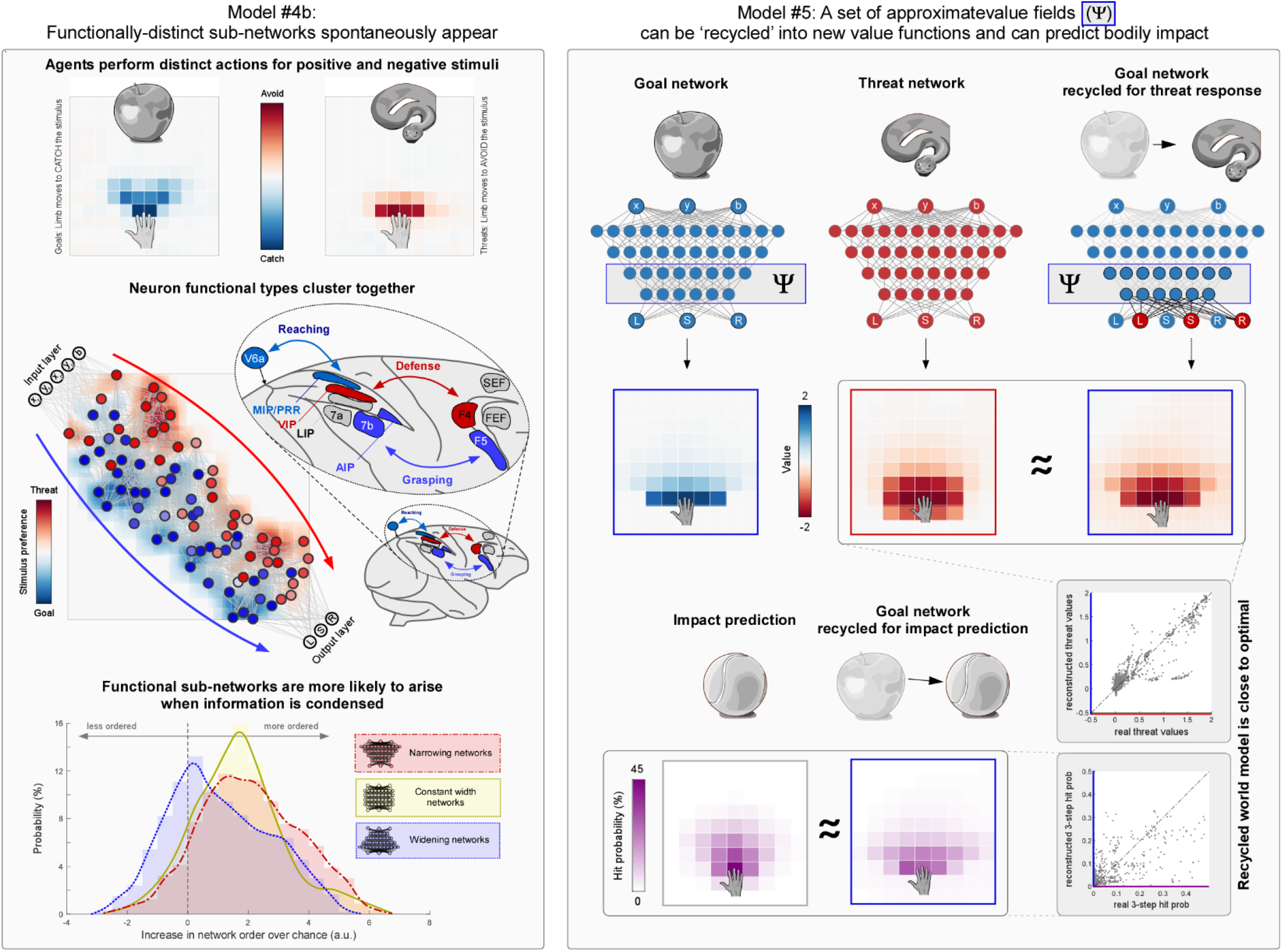

## Introduction

Many neural and behavioural responses have a spatial component. A class of such responses has offered a highly informative window into brain function: place and grid cells in the hippocampus and entorhinal cortex explicitly encode the animal’s position in its environment (1, 2).

Our functional understanding of place and grid cells has been enormously enriched by attempts to formally describe their activity patterns (1, 3–7). An example of particular relevance to this work came from the exchange of ideas between neuroscience and reinforcement learning (5), which demonstrated that *allocentric* place and grid responses – responses whose reference frame is anchored to the environment rather than the individual – are not purely spatial. Rather, they might provide a description of the world in terms of its future states (8, 9). This successor representation allows animals and artificial agents to interact with their environment at the spatio-temporal scale of locomotion and navigation to reach distant objects (5, 10). It follows that allocentric spatial responses provide an effective substrate for navigation in not only physical but also abstract, mental environments, even when facing a novel task (9, 11, 12). Subsequent empirical work has confirmed this prediction in humans and other animals (10, 13–15). Clearly, the substantial efforts to formally understand the computational mechanisms underlying *allocentric* responses have been enormously valuable.

However there exist other *egocentric* responses that have been less studied, both empirically and theoretically. Anchored to body parts, and dependent on the spatial *proximity* between stimuli and those body parts^1^, these responses have *peripersonal* fields (16–20). Peripersonal fields are measurable from macaque single neurons (21–25), human and non-human functional MRI (26–28), EEG signals (29, 30), and several behavioural measures^2^ (31–41). Such pervasiveness across scales and measures makes peripersonal responses excellent candidates for studying systems-level neuroscience. Nonetheless, while the theory behind place and grid cells is ever improving, formal theories of the function and necessity of peripersonal receptive fields are not yet as developed (42). Speculative notions have been put forward that these responses contribute to or reflect diverse and often vaguely-defined concepts such as body movement, the sense of self, impact prediction, and multisensory representation of space (19, 43–45). The few attempts at formalisation of peripersonal fields have largely focused on perceptual aspects (46–50), leaving several features unexplained. For example, existing models are often limited in both their temporal dynamics, and in the extent to which they are affected by motor repertoire and stimulus valence. Thus, a precise and unifying mathematical formalism reflecting a complete functional understanding of peripersonal fields is lacking (51).

Here we show analytically (i.e. normatively) and in-silico (i.e. computationally) that a reinforcement learning perspective can explain both the origins and the functions of peripersonal receptive fields. This perspective subsumes existing interpretations and formal models of peripersonal fields, and reproduces a range of foundational experiments. We also demonstrate that peripersonal fields provide a *short-term* map of the world *near* the agent in terms of its future states. This is analogous to how place and grid cells provide a *longer-term* map of the world at the *further* spatio-temporal scale of locomotion and navigation to reach distant objects. Together, these predictive maps, also known as successor representations, allow efficient interactions with a changing environment across multiple spatial and temporal scales.

## Results

### In theory: Peripersonal receptive fields can arise naturally from rewarded contact

Animals navigating the world experience reward or punishment upon contact between the body surface and objects in the environment. We contend that this simple fact fully accounts for the emergence of peripersonal fields centred around a given body part. We will first demonstrate this theoretically under a reinforcement learning framework, and then elaborate through computational modelling.

In a reinforcement learning model of an agent interacting with its environment, the values (*Q*) of actions that bring a body part into contact with certain objects correlate with the object’s proximity to that body part. When those *Q*-values are plotted as a function of stimulus position, graded fields surrounding the agent’s limbs arise (Figure 1). In this framework, three factors cause action values *Q* to correlate with proximity between objects and the agent’s body: (1) the temporal discount factor *γ*, (2) the world dynamics *p*(*s*_*t*+1_|*s*_*t*_), and (3) the actions *A* available to the agent (Box 1 and Figure 1). These three factors lead to the natural emergence of peripersonal receptive fields, as we explain below.

**Figure 1.**
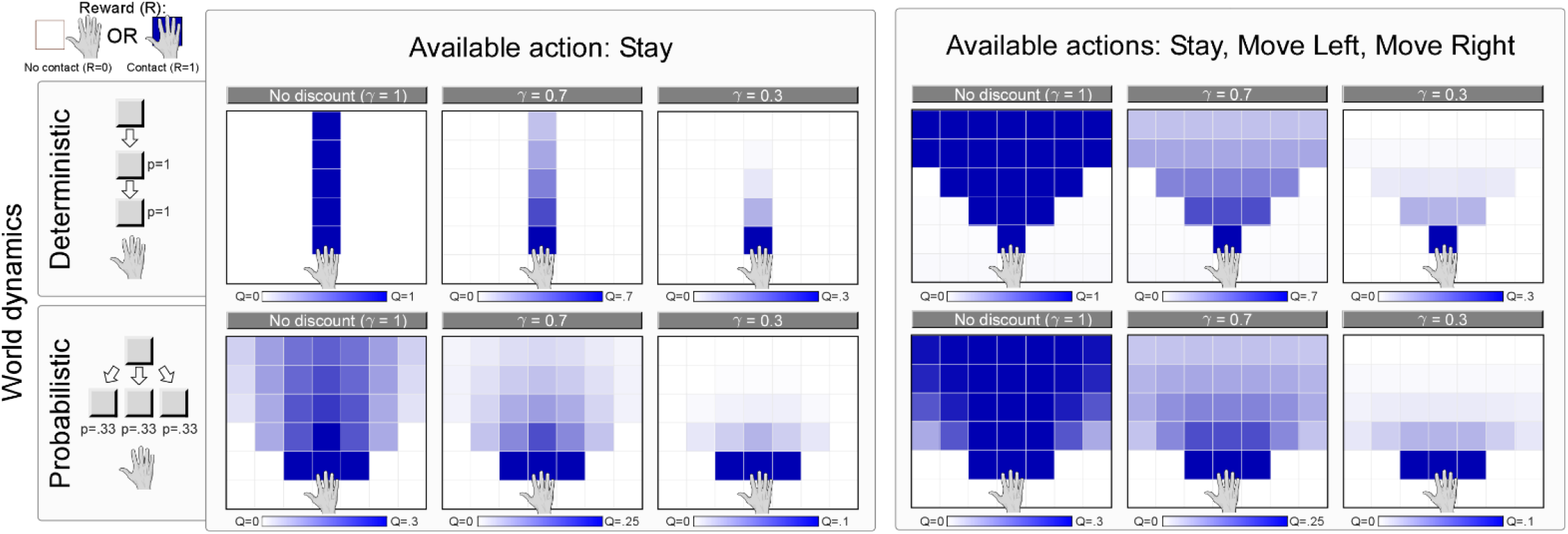
A reinforcement learning perspective explains the origin and properties of egocentric receptive fields. If contact between an object and a given body part is rewarded (top left; ‘Reward’), the value of an action that creates or avoids contact will form a receptive field centred around that body part. A number of additional factors determine the properties of such fields. 1) The world dynamics (rows) alter the probability that contact with a body part occurs, and affect how action value depends on proximity. 2) The actions available to an agent (main columns) also affect the positions from which an object can contact the body. Therefore, motor repertoire expands body-part centred receptive fields. 3) The discount factor gamma (sub columns) decrease the value of actions when objects are further from contacting the body in time. Thus, temporal discount γ creates an inverse relationship between stimulus distance to a bodypart and action value: a body-part centred receptive field.

#### 1) Proximity in time

The temporal discount factor *γ* causes the values of states *n* time-steps away from environmental contact with the body-part to be discounted by *γ*^*n*^ (Figure 1; Formally, 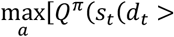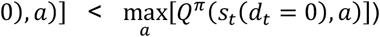. Given that spatial displacement of external objects takes time, the discount factor *γ* also causes the values of actions related to distant objects to be smaller than those related to objects closer to the body: the distance *d*_*t*_ is related to the number of timesteps *n* necessary for the object to contact the body. For example, if a wasp is near your hand you want to avoid the “negative reward”^3^ of a potential sting by moving the hand away. But if the wasp is still a few metres away from your hand, you do not feel an urge to either move the hand away or hit the wasp: the wasp is unlikely to your hand in the near future^4^. In this case, the number of timesteps *n* needed for the wasp to hit your hand is high. Therefore, actions related to avoiding or creating contact with the wasp are strongly discounted by *γ*^*n*^. In the simplest scenario, if the wasp (or any other object in the world) *always* moves towards the hand (i.e. *P*(*d*_*t*+1_ < *d*_*t*_) = 1), the value of actions should be monotonically related to the distance between the wasp and the hand *d*_*t*_ (Figure 1, top left trio of heatmaps). This relationship between value and distance naturally describes a body-part centric receptive field.

#### 2) Environmental dynamics

In a biologically more plausible case, the wasp is not guaranteed to move closer to the hand (i.e. *P*(*d*_*t*+1_ < *d*_*t*_) < 1). In this case contact is not certain to happen at any point in time, and more time will elapse until contact is made (if it is made at all). Therefore, the negative reward due to contact is weighted less strongly, and action value will fall off more strongly as a function of distance to an object *d*_*t*_ (Box 1; Figure 1, bottom left trio of heatmaps). This second factor further shapes peripersonal receptive fields, yielding maximal action value when the object is near a body-part, where probability of contact is highest. Even without any temporal discounting (i.e. *γ* = 1) action value will still be maximal near the body-part. Thus, environmental dynamics also contribute to the emergence of peripersonal receptive fields.

#### 3) Action repertoire

The repertoire of actions available to the agent also contributes to shaping peripersonal fields. Actions can alter either the distance to an object (*d*_*t*_; e.g., moving the hand further away from the wasp), or the probability that contact will be made (e.g., wearing a glove), or both. Because Q-values for *current* actions also take into account *potential* actions that can be made in the future (Box 1), it follows that Q-value fields can also appear in response to static stimuli: even if the wasp is dead, we can reach out to brush it away (Figure 1, right column).

Therefore, given that contact with objects in the environment is rewarded, action values naturally take the shape of peripersonal receptive fields.

### In practice: artificial peripersonal receptive fields strongly resemble biological ones

Despite its simplicity, the principle that object-body contact is rewarded or punished accounts for many behavioural and neurophysiological results. To demonstrate this, we trained a set of artificial neural network (ANN) agents to move one or two body parts (referred to onwards as ‘limbs’) in a 2D grid world (Figure 2). In a series of in-silico experiments, we recreated multiple foundational experimental findings describing peripersonal receptive fields. We obtained the following six results. (1) The agents’ behavioural responses, as well as the activity of many individual neurons in the underlying networks, show body-part centric receptive fields, which remain anchored to a body part when it moves. (2) The spatial extent of these peripersonal fields depends on both stimulus speed and direction relative to the body part. (3) These fields are altered by tool use, both plastically and dynamically. (4) They expand in response to stimuli of stronger valence. (5) In the ANNs underlying the agents’ behaviour, distinguishable sub-networks for goal-oriented vs threat-avoidance behaviour appear. (6) These ANNs provide a reliable statistical model of the environment near the agent, which can be recycled to respond effectively to novel tasks and environments.

**Figure 2.**
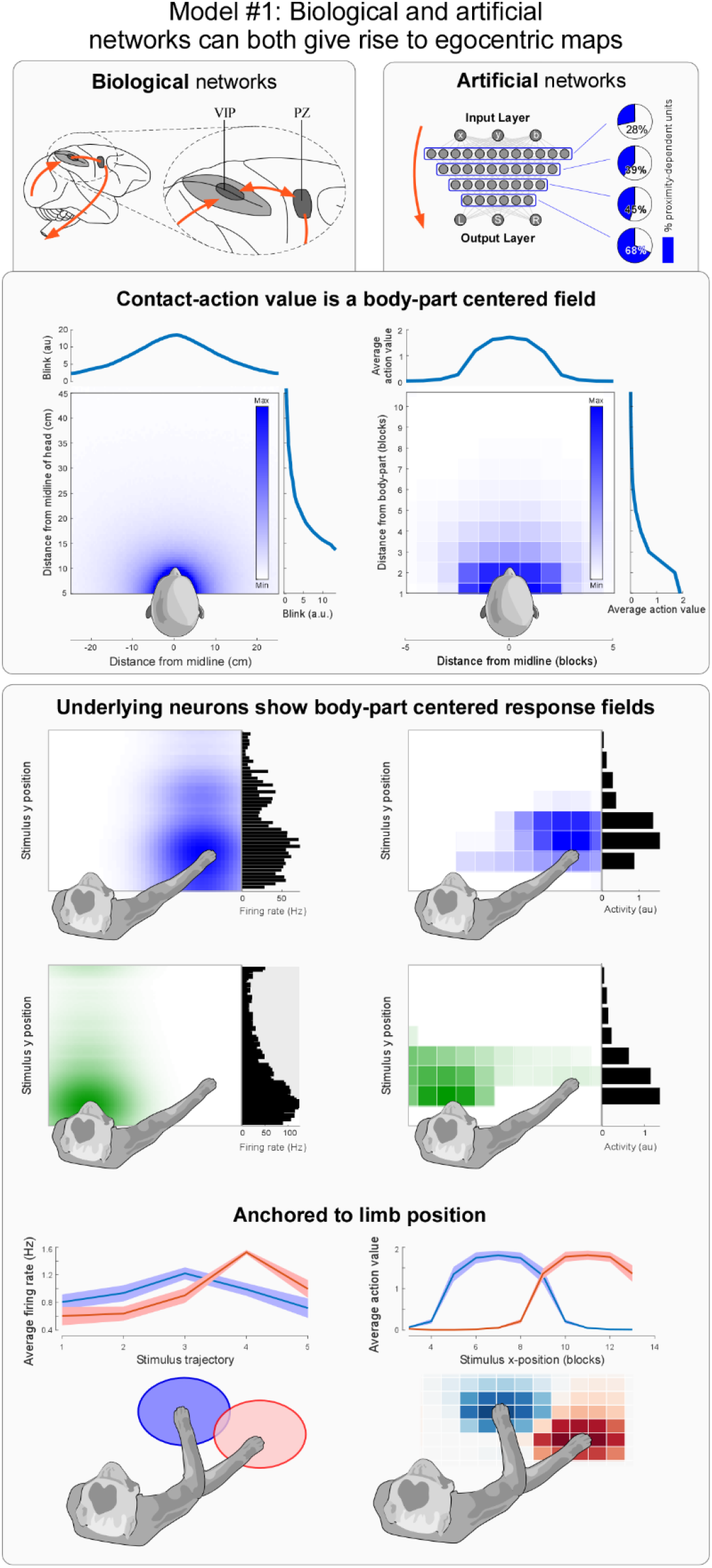
Artificial agents demonstrate peripersonal value fields. <u>Top panel:</u> Several behaviors that reflect the value of being hit by an external stimulus – e.g. the blink reflex – are graded as a function of stimulus distance to the face (left column). Artificial agents that receive reward or punishment after being hit by a stimulus show similar peripersonal fields reflecting their estimate of action value (right column). <u>Bottom panel:</u> Macaque brain areas VIP and PZ (left column) contain neurons with receptive fields around various body-parts (limb, 1^st^ row; head, 2^nd^ row; data reproduced from Graziano et al., 1994, 1997). Artificial networks (right column) contain similar neurons. Here we show responses from an agent trained to simultaneously move two ‘body-parts’. Different artificial neurons in the network (1^st^ and 2^nd^ row), respectively have ‘limb’ and ‘face’ centred receptive fields. The proportion of neurons with such receptive fields increases as a function of layer depth (top of right column). Peripersonal neurons in the macaque brain move their receptive field as a function of body-part location (left column, bottom row). The artificial neurons and resulting action values are also anchored to body-parts (right column, bottom row).

#### Box 1: Reinforcement Learning principles and model assumptions

In Reinforcement Learning, an agent attempts to maximise the cumulative future reward *R* by interacting with its environment through a set of possible actions *A*. Here we use a particular Reinforcement Learning technique, called Q-learning. The value *Q* of performing an action *a* in a state *s*_*t*_ at time *t* under a given policy *π* is the expectation E of the cumulative reward *R* discounted by *γ:*

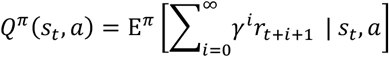

In order to maximise reward, the goal of Q-learning is to find an optimal policy *π*^∗^, which results in optimal action values 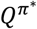 that satisfy the equation

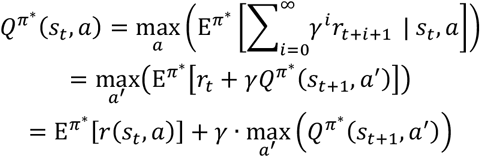

In other words, the optimal value of an action is the expected total discounted reward received when, after starting in a state *s*_*t*_, the agent performs an action *a*, and follows the optimal policy thereafter. To find these optimal action values 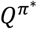, the agent iteratively updates its current estimate of value *Q*^*π*^: after the agent gains experience by interacting with its environment, the value of an action *a* in a given state *s*_*t*_ is updated as

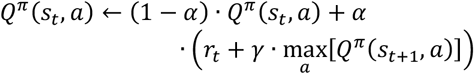

Where *α* ∈ [0,1] is the learning rate, *γ* ∈ [0,1] is the discount factor, and *r*_*t*_ is the reward. Over time, these updates bring *Q*^*π*^ progressively closer to the optimal value 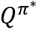.

We instantiate the assumption that contact between the agent and the environment is rewarded or punished, by dictating that when the distance between the body and an external object *d*_*t*_ becomes zero, *r*_*t*_ is non-zero, and (for simplicity) that *r*_*t*_ is zero otherwise:

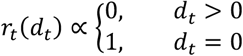

Under this assumption, the *Q* action values will manifest as peripersonal receptive fields, in which *Q* magnitude correlates with the object-body proximity.

##### Glossary

###### State *S*_*t*_

The configuration of the agent and the environment at a time *t*

###### Actions *A*

Movement options chosen by the agent at every timepoint. Different individual actions *a* may differently affect the state transition *s*_*t*_→*s*_*t*+1_ (i.e. the probability of reaching a specific state at the next timepoint)

###### Policy *π*

The method of choosing a single action *a* from all available actions *A*.

###### Reward *r*

An experience of the agent, occurring in certain states. The agent’s aim is to maximise summed reward over time. Reward can be either positive or negative.

###### Discount *γ*

The amount by which the agent de-emphasises the importance of future reward when summing predicted future reward

###### Value *Q*

The agent’s expectation of the total discounted reward *r* if it takes an action *a*, and thereafter follows a particular policy *π* to select between its future actions

###### *A*. Peripersonal field

When touch is rewarded, *Q* takes the shape of a body-part centric or *peripersonal* receptive field.

###### Task

The agent’s task is defined by the distribution of rewards across states. For example, in one task, an agent might experience a +1 reward upon contact with an object, while in another task the reward might be −2.

###### Successor features *ψ*_*t*_

A collection of predictive features that can be combined to form an effective model of the world: a **successor representation**. By performing a weighted sum over these features, a (new) action value *Q* can be calculated.

###### Peripersonal map

Individual successor features can simply be *Q* values for different tasks. Therefore **peripersonal fields** could be used as building blocks to form a successor representation of the world near the body: a peripersonal map.

#### Modelling Approach

Details about models are provided in supplementary methods. Briefly, in all models an agent is situated in a grid world. In this world, the agent can either keep its limbs still, or move them left or right. Objects appear at the top of the grid world and generally move along a downward trajectory towards the agent, with some random noise at each timestep. The exact speeds and trajectories differ slightly between models (see Methods for details). An object can either be a ‘goal’ or a ‘threat.’ Agent contact with a goal results in a positive reward, and agent contact with a threat results in a negative reward. For each model we tested multiple ANN architectures to statistically test consistency of experimental results across different network types (see Methods for further rationale) This approach deliberately abstracts over specific physiological details, demonstrating that the results obtained in biological systems can be explained from a set of simple basic assumption and through a simple mechanism. The following sections describe each model in detail.

#### Model #1 – Artificial agents produce response fields anchored to the limb

##### Background

A defining feature of peripersonal fields is that response magnitude (of either behaviour or single neurons) correlates with the distance between environmental stimuli and the body (Figure 2, right column). Crucially, this correlation is invariant to body-part position, i.e. it does not change when the limb is displaced (Figure 2, bottom row). This observation, which has been demonstrated in a plethora of studies using both visual and auditory stimuli, has often been interpreted as a coding of stimulus position in egocentric coordinates (16, 17, 20, 52–56).

Neurons showing such proximity coding have been described in the putamen and the parieto-premotor system, most prominently in ventral intraparietal area (VIP), and the dorsal part of F4, near the boundary with F5 (21–25) (Figure 2 top left panel). These neurons are often referred to as peripersonal neurons, and many of them also respond to somatosensory stimuli delivered to the body-part around which the visual or auditory receptive field is centred.

Behavioural measures with similar response features have been identified in humans, typically using concomitant audio-tactile and visuo-tactile stimuli (57, 58). Several imaging studies have shown overlap between human and macaque brain areas which contain or modulate peripersonal fields (27, 59–61).

##### Approach

We designed Model #1 to test whether similar *artificial* limb-centric response fields would emerge from action value. We ran this using 6 different network architectures. In 3 of these 6 instances the agent controlled 1 limb, and in the other 3 the agent controlled 2 limbs. The environment only contained goals. For more details, see ‘Base Model’ and ‘Model #1’ in Methods.

##### Results

The action values in all architectures of Model #1 were highly reminiscent of the biological responses: they consistently correlated with proximity between the object and the agent’s limbs (p ≤ 7.97×10^−3^, |ρ| >0.086, FDR-corrected Pearson correlation tests; Figure 2). Similarly, a substantial proportion of units within the networks showed receptive fields with body-part centric distributions, regardless of exact network architecture and the number of limbs controlled by the agent (51 ±9% [SD; from here on out all Figures following ± will be standard deviation SD]; Figure 2; see supplementary methods for the precise network architectures tested and the rationale behind them). This was especially the case in the later network layers, where units were closer to representing action value (p = 1.56 ×10^−9^, LME main effect of layer depth; 63 ± 21% of units being body-part centric in the last layer; Figure 2).

##### Conclusion

These results indicate that *biological* neurons which show peripersonal receptive fields anchored to the limb might *also* be outputting the value of an action which is rewarded by contact with a stimulus, as we have suggested elsewhere (16).

#### Model #2 – Artificial peripersonal fields are reshaped by stimulus dynamics

##### Background

A second property of almost all biological peripersonal receptive fields is that in they are also altered by many non-spatial variables, similarly to place cells (14, 62).

For example, peripersonal fields – whether neurophysiological or behavioural – expand when the velocity of a looming stimulus increases (63–66). They are also often anisotropic, with a maximal extent in the direction of movement of the stimulus relative to the body-part (24, 65, 67). In theory, action values of an artificial agent should similarly depend on stimulus velocity. They should depend on speed because a faster approaching stimulus can travel more distance between timesteps. They should also depend on direction, which dictates whether the stimulus will hit the body.

##### Approach

To demonstrate these kinematic dependencies, in Model #2 we exposed 4 agents (each subserved by a different ANN architecture) to stimuli moving at different speeds and in different directions. For details, see ‘Model #2’ in Methods.

##### Results

As predicted, action-value fields expanded when stimuli moved faster (p≤8.46×10^−3^, ρ ≥0.42, FDR-corrected Pearson correlation tests Figure 3, top), and specifically in the direction from which stimuli approached (p ≤3.24×10^−12^, ρ≥0.85, FDR-corrected Pearson correlation tests; Figure 3, top).

**Figure 3.**
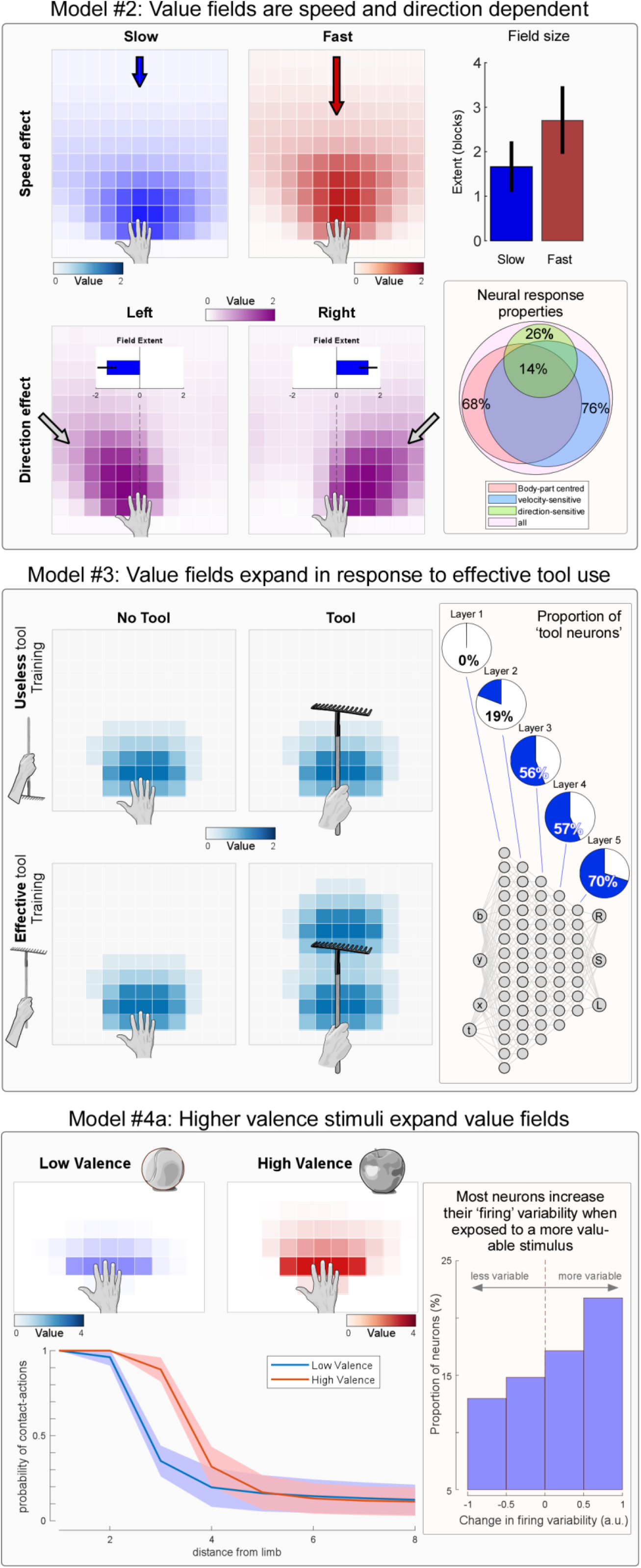
Artificial peripersonal fields demonstrate most of the properties of biological peripersonal responses. <u>Top panel:</u> Canonical biological peripersonal fields depend on stimulus velocity and direction. Artificial value fields also expand when incoming stimuli move faster (top row), in the direction of the incoming stimuli (bottom row). A substantial proportion of the neurons in the agent’s network are velocity sensitive (i.e. speed and direction; right column). <u>Middle panel:</u> Canonical peripersonal fields extend to incorporate the tip of a tool, but only after training with the tool. Artificial value fields also expand only after training with a tool that increases the ability to touch an object (bottom right sub panel). Artificial neurons also have receptive fields that expand as a function of tool use after training. These neurons are prevalent in deeper network layers (right column). <u>Bottom panel:</u> Canonical peripersonal fields are larger and have greater magnitude in response to stimuli of higher valence. Artificial value fields show similar properties (1^st^ and 2^nd^ columns). This means that contact creating or avoiding actions are initiated when a high-valence object is at a further distance from the body (bottom graph). Individual artificial neurons are also more affected by the location of a high-valence than a low-valence stimulus (3^rd^ column). This reflects the biological observation that the brain regions containing peripersonal neurons show substantially stronger activity to behaviourally relevant stimuli (77, 126).

Activity of individual artificial neurons conformed with these observations. The extent of individual receptive fields correlated with stimulus speed and direction (26 ± 6% of units had receptive fields whose extent correlated with vertical movement speed; 76 ± 8% expanded in the horizontal direction of movement; 20 ± 6% correlated with both; Figure 3, middle panel).

##### Conclusion

Such mixed encoding of position and velocity parameters is another physiological hallmark of peripersonal neurons, and of the brain areas in which they are found (17). These results strengthen the case that biological peripersonal neurons represent late stages in a system computing contact action value.

#### Model #3 – Tool use reshapes artificial peripersonal fields

##### Background

During tool use, certain peripersonal neurons expand their receptive fields to incorporate the tool and the area surrounding it (68). Other behavioural and neural measures are similarly affected (69–73). This plastic remapping of receptive fields^5^ seems to only occur after some initial experience with the tool: if participants – whether human or monkey – hold the tool without having used it, no change in receptive fields occurs. People with extensive tool use, however, are different. For example, as soon as blind cane users pick up a cane, their hand centric response fields dynamically change (74).

##### Approach

To demonstrate these remapping effects, in Model #3, we first trained 3 agents that occasionally held a ‘tool’^6^, but the tool was *ineffective*: stimulus contact with the tool tip did not reward the agents (see Model #4 in Supplementary Methods). Subsequently, we re-trained the same agents while they held an *effective* tool that, when touched by the object, granted reward to the agent.

##### Results

We observed a main effect of training, a main effect of tool presence, and an interaction (2×2 ANOVA; p = 1.37×10^−6^, p = 6.37×10^−26^, p = 1.08×10^−6^ respectively). To investigate the source of these effects, we performed post-hoc comparisons between conditions. As expected, without effective tool training there was no evidence of remapping when picking up the tool: there was no consistent effect on the Q-fields between conditions in which the agents did or did not hold the tool (1.0 ± 0.0 vs 1.1 ± 0.3 Q peaks; p = 0.125; Wilcoxon Signed Rank test). However, after training the agents while they held an *effective* tool, the presence or absence of the tool strongly affected the Q-fields (1.4 ± 0.5 vs 2.0 ± 0.0 Q peaks; p = 2.73×10^−6^; Wilcoxon Signed Rank test). This was due to the appearance of a new Q-value maximum around at the tip of the tool, but only when the agent held the tool (Figure 3, middle). Therefore, after training the receptive fields had *plastically* adapted, and *gained the ability to dynamically* adapt Q-values depending on tool presence. A substantial proportion of units within the networks showed the same type of variation in their receptive fields as a function of tool use (50 ± 5%; Figure 3). This was especially true in later network layers, where units were closer to representing action value (p = 4.34 ×10^−5^, LME main effect of layer depth; 70 ± 22% of units displaying receptive field remapping; Figure 3, middle).

##### Conclusion

Therefore, both plastic and dynamic changes due to tool use are consistent with action value coding. Our model shows that plastic changes in peripersonal responses might represent long-term learning processes involving changes of neural connectivity, while dynamic changes might instead represent pre-learned associations that are simply instantiated through differences in neural activation in different states. Such an interpretation does not require the common assumption that a change of “body schema” is necessary to explain tool use remapping (75, 76).

#### Model #4 – Valence alters peripersonal fields and underlying networks

In Model #4 we describe the effects of magnitude and sign of rewards. We show that these factors respectively contribute to the expansion of peripersonal fields, and the neuroanatomical segregation of movement networks.

##### Artificial peripersonal fields expand when exposed to high valence stimuli (4a)

###### Background

Stimuli with higher valence (e.g., highly desirable or highly dangerous objects) elicit stronger responses from biological peripersonal neurons and brain regions that contain such neurons (77–79)^7^.

###### Approach

To investigate whether the action value framework instantiated in our models could recapitulate this known effect of stimulus valence on response fields, we varied the value of the reward *r*_*t*_. We predicted that a larger absolute reward would lead to overall higher action values. To test this prediction, we trained 3 agents to respond to stimuli of various reward magnitudes.

###### Results & Conclusions

All agents showed larger response magnitudes for higher reward stimuli (p ≤ 2.42×10^−67^, |Z| ≥ 17.3, Signed Rank test). Consequently, actions related to a higher reward stimulus were performed at greater distances from the body (p = 9.77 ×10^−4^, FDR-corrected Signed Rank tests; Figure 4, bottom). This mirrors behavioural studies, which describe a response field expansion when stimulus valence is higher (34, 80–83). In fact, even conditioned fear responses seem to be learned in a body-part centric manner (84).

**Figure 4.**
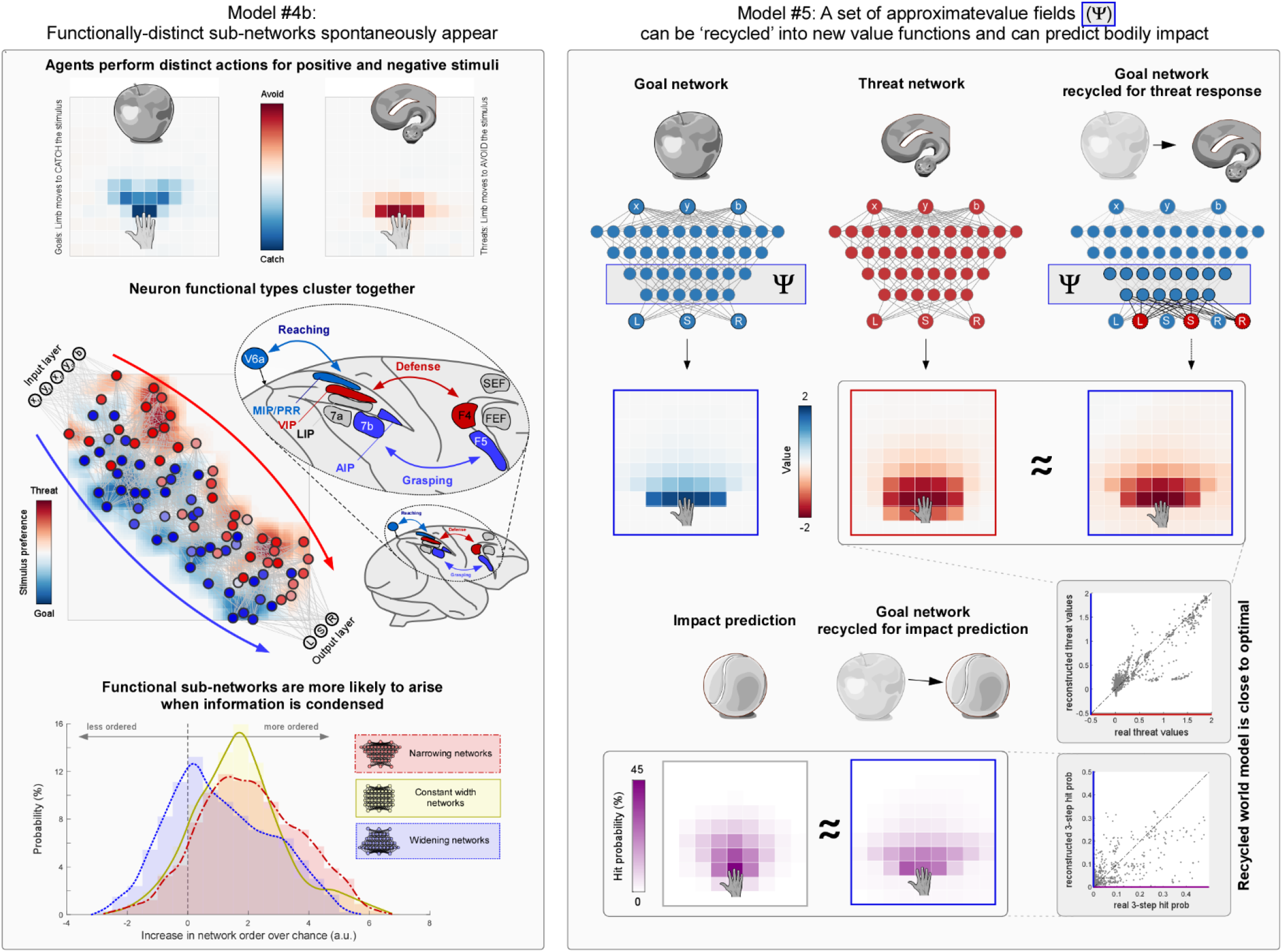
Network analysis: Functional subnetworks and Successor-representation world models. <u>Left panel:</u> Motor areas in the primate brain contain distinct functional units that coordinate different types of behavior. Similarly, training an artificial network to perform both approach and avoidance behaviors (top row) gives rise to spatially distinguishable sub-networks (middle row). Individual neurons can be classified as threat- or goal-preferring (red and blue, respectively). When the neuron-to-neuron distance depends on connection strength, identifiable threat- or goal-preferring sub-networks appear (red and blue, respectively). This is reminiscent of the anatomical structure of the parieto-premotor system, where peripersonal neurons cluster together based on their behavioural function (inset). Such sub-network structure is particularly likely to appear when the network condenses information (i.e. when it narrows; pink histogram), compared to when it spreads out information over many neurons in later layers (i.e. when it widens; blue histogram). <u>Right panel:</u> Peripersonal fields could be used as basis functions to flexibly interact with the world near the body. An artificial network that has only learned to reach positive valence stimuli (left column, blue; top row) can be ‘recycled’ to approximate an appropriate value field for avoidance movements (center column, red; top row). Specifically, by taking a weighted sum of the neural activities in the second half of the blue network (ψ), the output from the red network could be faithfully reconstructed (last column, red fields). Furthermore, the probability that a stimulus would hit the body over any number of timesteps (3-timestep hit-probability shown; left purple field, bottom row) could be faithfully reconstructed using the same second half of the blue network (*ψ*). This is particularly informative given that the agent never had access to information more than 1 timestep back, while the derived hit-probability is for 3 timesteps in the future: action values allow the agent to build up a longer-term predictive model.

Individual artificial neurons were also affected by reward magnitude. When the networks were exposed to high value stimuli (i.e. with higher absolute *r*_*t*_), neural activity varied more than when exposed to low-value stimuli. (p = 2×10^−3^, Z =2.738, Signed Rank test). This mirrors the fact that brain regions including peripersonal neurons respond more strongly to high-valence stimuli than to motivationally irrelevant ones (77, 78).

##### Valence sign results in distinct sub-networks subserving different movement classes (4b)

###### Background

Peripersonal responses are often said to be related to separate classes of movements. Specifically, a distinction is frequently made between defensive avoidance movements, and grabbing or appetitive movements (17, 85–87). This distinction is based on the type of movements associated with the brain areas where peripersonal neurons are found: VIP and F4, for instance, are more linked to defensive movements, while MIP, AIP, F5 and 7B are more related to reaching and grasping (86). In fact, microstimulation of peripersonal VIP neurons elicits defensive movements. Similarly, biasing firing rates in the dorsal part of F4 using bicuculine and muscimol alters both likelihood and magnitude of defensive responses (88). Unfortunately, no other neurons with body-part centric receptive fields have been directly stimulated in other brain areas. Still, prolonged electrical stimulation of large areas of pre-central cortex results in spatially-specific movements that often seem to be appetitive (89).

It remains debated why these action classes are represented in different cortical areas, and how they relate to peripersonal responses (52, 56, 85, 87, 90). Our model offers a perspective that clarifies the issue: the separation between classes of actions and their subserving functional networks can be understood by considering the effect of the *sign* of the reward *r*_*t*_, which is equivalent to stimulus valence.

###### Box 2: Successor features

Successor features (*ψ*^*π*^) are a generalisation of the successor representation which has gained traction in both AI and neuroscience (11, 12). Successor features provide a possible solution to the problem of relearning value functions. The value function 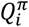 for a specific task *i* and a policy *π* can be constructed from two parts: the successor features *ψ*^*π*^ and the successor weights ***w***_*i*_.

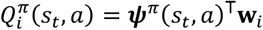

This allows the same successor features *ψ*^*π*^ to be re-used for different tasks *i* by simply using different weights *w*_*i*_. Place cells probably provide successor features for a successor representation with temporal and spatial scales that allows animals to navigate through the environment (9, 11). This fits with the hippocampus’ role as a coordinator of multiple regions across extended periods of time (127).

In the main text we demonstrate that peripersonal fields might provide successor features analogous to place cells but at smaller spatial and temporal scales. Similarly to the way in which place and grid cells allow navigation through the distant environment, body-part centric receptive fields might allow interaction with the environment near the body.

This notion of a peripersonal, egocentric *map* as a near-body world model is supported by a recently demonstrated advantage of successor features: they can be approximated by a collection of value function estimates 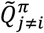 across multiple tasks (128):

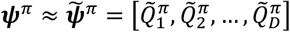

Thus, a peripersonal value function for a novel context *i* should simply be computable as a linear combination of several other peripersonal value functions (or, more realistically, of the activity of multiple late-stage units approximating those value functions; see right panel in Figure 5).

A further demonstration of value fields as world models is the calculation of impact prediction. Impact prediction is a particularly easy world model to compute using peripersonal value fields, given that 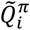 is a function of the probabilities of bodily impact, under task *i*, conditioned on action policy *p*(*s*_*t*+*k*_ = touch|*s*_*t*_ = vision or audition, *π*) (see Box 1):

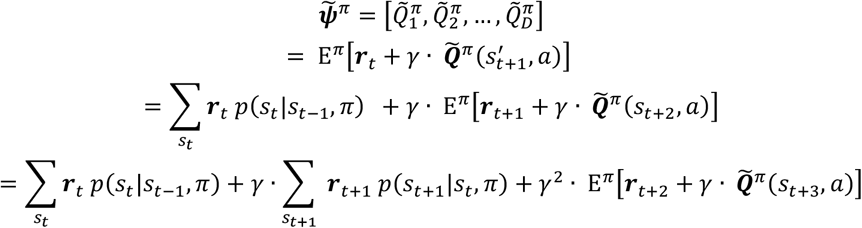

etc. Then, given that 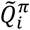 is the value function *for creating or avoiding bodily contact* (i.e. only touch is rewarded), we can set ***r*** to zero if no touch occurs, and the contribution of contact probability becomes explicit:

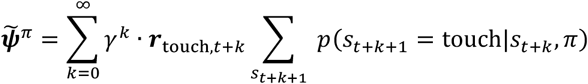

**Figure 5.**
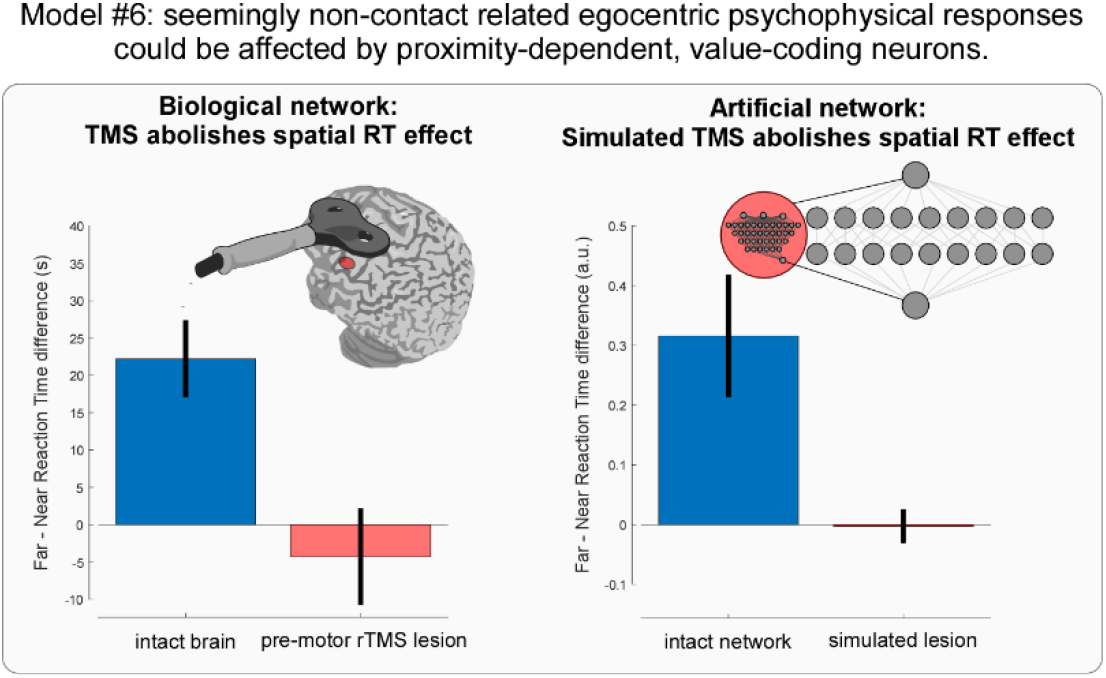
Human reaction times to tactile stimuli shorten when concomitant auditory or visual stimuli are near the body. This effect is abolished when posterior parietal cortex or ventral pre-motor cortex is reversibly lesioned with TMS (left column; data from (59)). Similar results can be obtained for an artificial agent. If the agent’s network contains a module (red circle) that calculates contact-action value, reaction times to tactile stimuli will be shortened with visual stimulus proximity. When the contact-value module within the main network is ‘lesioned’, this effect vanishes (right column).

###### Approach

To investigate this possibility, we exposed 45 agents to two types of objects (the rationale for the number of agents is explained in Model #4b of Methods). One type (goals) led to positive reward upon contact with a limb, while the other (threats) led to negative reward (i.e. a punishment; see footnote 1 above).

###### Results

Both types of objects resulted in canonical action-value fields graded with limb proximity and anchored to limb position (p ≤ 8.98 ×10^−4^, ρ ≤ 0.124 for positive valence stimuli; p ≤ 5.59 ×10^−6^, ρ ≥ 0.163 for negative valence stimuli; FDR-corrected Pearson correlation tests; Figure 4, left). Behaviourally, the agents moved their limb towards positive objects, and away from negative objects (p ≤ 5.82 ×10^−4^, |Z| ≥ 3.44; Figure 4, left).

The activity of substantial proportions of units within each network correlated with proximity of at least one stimulus type (79 ± 7%). Units showed a clear preference for different stimulus types: 60 ± 12% of units’ activities correlated with proximity of goals, 55 ± 12% with the proximity of threats, and 36 ± 10% with both. Interestingly, in almost every network instantiation, two spatially-segregated sub-networks reminiscent of those described above in primates emerged: positive-valence preferring units were more strongly connected to each other, and vice-versa (p ≤ 0.05 for 32/45 networks, FDR-corrected Pearson correlation tests). As a result, there was more sub-network structure than expected by chance across all networks (p ≤ 5.59 ×10^−6^, T = 7.29; un-paired t-test; Figure 5).

###### Conclusion

Thus, the anatomical separation of the cerebral cortex into multiple action classes (91, 92) could be an emergent property of the different requirements of actions afforded by different types of stimuli: these actions are more efficiently deployed if the networks processing the stimuli separate at early stages. Should the networks not segregate at early stages, the necessary differential processing resulting in different action types would be left to the final stage of motor transformation. In a system with limited bandwith this would result in a lower SNR than necessary (93, 94).

### Peripersonal action values as near-body world models

#### Model #5 – Transferable Successor Features

##### Peripersonal neurons can be recycled to perform novel tasks (5a)

###### Background

Context and environment can change quickly. Thus, both biological and artificial agents are more effective when they are quickly able to update their behaviour upon encountering a novel task or environment. For novel tasks, recomputing action values from scratch is highly inefficient and can overwrite previously learnt associations (95, 96). If the agent instead has access to a predictive model of the world, it can simply multiply the probability of future states by the expected rewards, and thereby generate appropriate action values to novel situations (96, 97).

We will now demonstrate that this principle extends to peripersonal value fields: they can be used as building blocks that are recombined into novel action values. From this perspective, peripersonal receptive fields are successor features (see Glossary), reminiscent of place cells in hippocampal function (9) but working at shorter spatio-temporal scales (see Box 2). Thus, a sufficiently large number of peripersonal value fields create a model of the world near the body, an *egocentric map* that allows agents to cope with an ever-changing environment.

###### Approach & Results

To demonstrate that peripersonal receptive fields can be used as transferable successor features, we computed new value fields for novel stimuli to which network A had never been exposed. Specifically, we first trained network A by only exposing it to positive stimuli, and network B by only exposing it to negative stimuli. We then approximated the value functions from network B (i.e., value functions appropriate for responding to threats) by using a linear combination of value functions coming from network A, which had never been exposed to threats.

Note that this approximation is not trivial for two reasons. First, the value field for a given action in the presence of a goal is not simply the inverse of its value when there is a threat^8^. Second, the dynamics of the environment that networks A and B were trained in were different, to make reconstructing a negative-valence value field harder and thus demonstrate generalizability across environments. Specifically, instead of a 50% chance of moving an additional block up, down, left or right (i.e., the general rule of object movement in all other models; see Methods), goals had a 33% chance of moving an additional block left or right, and never moved an additional block up or down. Nonetheless, the linear combinations of goal value functions calculated by network A were excellent approximations of the threat value functions computed by network B (p ≤ 10^−30^, ρ ≥ 0.87, FDR-corrected Pearson correlation tests).

Similarly, we recreated the value functions of network B using only the activity of units in late layers of network A (i.e., units in layers close to the output layer; those computing positive reward-seeking values). These threat-avoidance values were good approximations of the originals (p ≤ 10^−30^, ρ ≥ 0.83, FDR-corrected Pearson correlation tests p ≤ 10^−30^, ρ = 0.92, FDR-corrected Pearson correlation tests; Figure 4, right panel). Therefore, value functions that had been learned in order to reach an appetitive stimulus, can be easily recycled to avoid an aversive stimulus. Crucially, such a mechanism *avoids having to completely relearn threat-specific value functions from scratch*.

###### Conclusion

This recycling of value functions is a practical example of how a set of peripersonal fields could constitute the building blocks for a successor representation of the world near the body (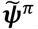; Box 2 and Glossary). The successor representation is recycled to allow the agent to interact adaptively with the near-body environment. Such a flexible, value-based world model provides a formal explanation for the notion that the representation (and even perception) of the space near the body is built up of a set of motor schemata (17, 53, 98, 99). Our formal explanation is also fully in line with the conceptualisation of posterior parietal areas as state-estimators (100) that use partially overlapping motor codes (101).

##### Subsuming other formal explanations of peripersonal responses (5b)

###### Background

There are other interpretations of the functional significance of peripersonal responses. Many interpretations postulate that these responses reflect two types of world models: *impact prediction* and *multisensory integration* (46, 47, 49, 102–104). In fact, both models entail the same computation: both use vision or audition to compute the probability of touch (i.e. 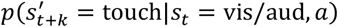, but the prediction occurs at different timescales (51). In multisensory integration touch is predicted in a short time-window around the visual or auditory stimulus, while in impact prediction, the computation occurs across a longer time-window, which always extends forward in time from the visual or auditory stimulus.

Our successor representation subsumes and accommodates these previous explanations into a more general formalism. It explains, for example, why one can predict impact from peripersonal responses: value functions resulting from rewarded touch include terms which are essentially discounted estimates of hit probability, conditioned on action choice policy and multiplied by the value of contact reward (see Boxes 1 and 2).

###### Approach & Results

It follows that action values (as well as the activity of units in late network layers) can be used to approximate the probability that a stimulus contacts the agent’s body at any point in the future. To demonstrate this, we approximated hit probability 1 to 7 timesteps into the future, using a linear combination of value functions coming from network A (see model 5a). These linear combinations were excellent approximations of hit probability at all 7 timespans (p ≤ 10^−30^, ρ ≥ 0.63 using action values, p ≤ 10^−30^, ρ ≥ 0.78 using neural activity, FDR-corrected Pearson correlation tests; Figure 4).

###### Conclusion

In other words, the ability to predict touch on the basis of visual or auditory input is an arising property of the peripersonal successor representation. Thus, this perspective subsumes existing formal explanations based on impact prediction and multisensory integration.

A further advantage of the formalism we propose is that most other formal models require highly specific assumptions about what the brain might be encoding or about the underlying neural architecture. For example, some older models of peripersonal responses specifically assumed that the brain encodes multisensory integration, thus forcing the network architecture to combine visual and tactile inputs by fixing specific weights between specific units (50). In other words, the inevitable result that a certain model successfully performs, say, function A (e.g. multisensory integration) when that model is set up to do precisely A, does not tell us why function A is achieved by the brain. Newer models used a lighter touch with respect to network architecture, and were able to recreate many features of canonical peripersonal space responses. However, they did still explicitly assume that the brain encodes impact prediction (47, 49, 104, 105), multisensory integration (106), or limb-centric target position (107) without providing *a priori* reasons for why the brain might do this. In a recent neural network model, Bertoni et al (46) showed that peripersonal fields can arise under the more parsimonious assumption that the brain attempts to condense multisensory input information. While this result is interesting, such pure representational models inevitably miss out the main function of the brain: acting to survive (93, 108–112).

In contrast, here we started *a priori* from the basic assumption that the brain acts to survive by maximising reward and minimising punishment through making or avoiding contact with objects in the environment. We subsequently demonstrated that the abilities to perform multisensory integration and predict impact are natural consequences of computing action value, without having to assume that they are being encoded *explicitly* by the brain. We also showed that impact prediction is conditional on the available actions (Figure 1), something which impact prediction models, as well as other purely representational models tend to ignore^9^.

#### Model #6 – Artificial psychophysical responses that seem unrelated to creating contact nonetheless form peripersonal fields

##### Background

A number of behavioural responses that on the surface seem unrelated to creating contact are nonetheless proximity-dependent. For example, tactile *detection* reported by button press – i.e. an action that does not create contact with the concomitant auditory or visual stimulus – improves when the concomitant stimulus is delivered near the tactile stimulus (34–36, 113). This proximity-dependence can be smoothly explained by the successor representation perspective.

The reason is simple. Stimuli occurring at certain distances from the body, with or without an explicit instruction to grab or avoid them, activate a network that reflects the value of *potential* grabbing or avoidance actions: the peripersonal successor representation. This activation facilitates motor output for all actions that have overlapping motor codes with the successor representation. Given that an untrained and novel task involving near-body responses should make use of the successor representation, it follows that the tactile detection task, will be facilitated by the peripersonal successor representation, and thus show proximity dependence. Notably, tactile detection stops being affected by the proximity of the non-somatosensory stimulus after rTMS lesions of brain regions strongly related to motor planning and containing peripersonal neurons (e.g. posterior parietal and ventral premotor cortices) (114).

##### Approach

Here we demonstrate the above postulate: behaviours that do not lead to creating or avoiding contact could nonetheless show proximity dependence because they are affected by proximity-dependent neurons that calculate action value. We trained a neural network to perform a new ‘reaction time’ task. In this task, the agent was required to report whether it received somatosensory stimulation, which was delivered concomitantly to task-irrelevant visual stimuli presented at different distances from the limb (see Model #6 in Methods). Crucially, the network contained a pre-trained subnetwork module that returned the Q values of moving under the assumption t hat contact with visual stimuli would result in a reward (Figure 5; module is described in Model #1 of Methods). In other words, this ‘reaction time’ network made use of a peripersonal map: it included neurons that were also part of a ‘body-part centric response field’ network. For more details, see Model #6 in Methods.

###### Box 3: Model predictions

The formalism described here makes a number of empirically testable predictions.

**First**, extensive training should reduce or remove the proximity-dependent effect of a visual or auditory stimulus on the detection of a concomitant tactile stimulus. The reason for this prediction is that – under our framework – the network responding to the tactile stimulus uses a successor representation (i.e. the network overlaps with existing action representations meant to create or avoid contact; Model # 6). Therefore, after extensive training with feedback in a tactile reaction time task, the network enacting the task should progressively separate from the contact-sensitive network. Because the contact network responds to auditory and visual stimuli, the modulatory effect of such stimuli should lessen with training.

**Second**, training participants to better predict whether or when a stimulus will impact the body, should *not* affect peripersonal fields. The reasoning here is simple: impact prediction should be an arising property of action value, and thus computable *from* it. Hence, improving impact prediction should be equivalent to training the network that extracts information from the successor representation, and as such leave body-part centered fields unaffected.

**Third**, the extent of receptive fields should depend on certain traits that have analogues in reinforcement learning. One promising variable for investigation is the discount factor. Participants – human or animal – that show signs of being impulsive, can be said to have strong temporal discounting. As such, we should expect the receptive fields of these individuals to extend less far in both space and time. Similarly, one might expect that the state of an animal would affect its impulsivity and temporal discounting: if an animal is particularly hungry, it should be willing to wait longer and work more for food. As such, its receptive fields for appetitive actions should expand. The temporal discount factor could even be influenced by changing stimulus dynamics. Indeed, individual traits such as anxiety and cynophobia (fear of dogs) have been shown to affect peripersonal fields (129, 130).

**Fourth**, the effect of uncertainty in stimulus position and direction should be two-fold. On the one hand, increasing speed and position uncertainty should increase the zone of the receptive field with magnitude above a given cut-off, as it becomes more likely that stimuli might reach the agent from formerly unlikely positions (Figure 1). On the other hand, the maximal response magnitude within the receptive field should decrease: if the agent is less certain that a stimulus, even if very near, will move in its direction, the value of avoiding or catching it will necessarily decrease (Figure 1).

**Fifth**, decreasing the uncertainty of stimulus position and velocity to 0 should *not* homogenise peipersonal fields *as long as the agent is able to move and reach the stimulus*. This prediction is particularly significant because it runs counter to most other formulations of peripersonal fields. In those formulations that rely purely on impact prediction, it is exactly the uncertainty that creates the receptive field, because a stimulus that is immobile has no probability of contacting the body (46, 131). In our formalism however, the value of moving in response to a completely predictable and stationary stimulus is still non-zero, giving rise to a body-part centred receptive field (Figure 1).

**Sixth**, a participant or animal’s available motor repertoire should affect peripersonal fields. For example, immobilising a limb alters the available actions, and should thus affect action values, even for actions which have not been directly impeded (moving the arm – for example – will be much less valuable if the hand is immobilized than if it is free to grasp). Indeed, recent empirical results suggest this motor repertoire affects peripersonal responses (132, 133). Similarly, the actions afforded by a stimulus (29, 38) or one’s own limb dynamics (134) seem to affect receptive fields.

We note that the outcome of these predictions should not be used be used as proof that the proposed framework is entirely wrong or right. More likely, different measures will conform to these predictions to different extents, and should thereby help us understand to what extent a ‘successor representation with action-field components’ is a satisfactory explanation of the data at hand. If the prediction is entirely wrong, we need to supplement or replace our model for that measure. If the prediction is entirely right, we must keep searching for other exceptions proving that the framework is unsatisfactory.

##### Results

Reaction times to tactile stimuli were affected by the position of a concomitant visual stimulus, with shorter responses when the visual stimulus was near the limb (p=0.0024, Wilcoxon signed rank test). However, when the network module with peripersonal response fields was artificially lesioned, the proximity effect disappeared (p=0.0117, Z =2.5219, Wilcoxon rank sum test between lesioned and non-lesioned reaction times; Figure 5).

##### Conclusion

Together with TMS observations in humans (114), these results illustrate that when a network that calculates contact-action value (i.e. the peripersonal map in posterior parietal and ventral premotor cortices) is part of a larger motor network (91) that performs a behavioural task unrelated to creating contact, the task can still be modulated in a body-part centric fashion. Therefore, behavioural measures such as a tactile detection with concomitant task-irrelevant stimuli can be interpreted as a generalised index of multiple action values (16): the peripersonal map that affects these measures consists of multiple peripersonal fields describing the value of various contact-related actions. Therefore, these behavioural measures modulated by body-part proximity likely reflect an arbitrarily weighted average across multiple action values.

## Conclusion

We have demonstrated that body-part centric response fields are naturally arising properties of two simple and plausible assumptions about living agents. First, agents experience reward or punishment upon contact between the body and external objects. Second, they act to maximise reward.

The resulting receptive fields can be observed both in artificial single-neuron activity and in the behaviour of the agent. These artificial fields are sensitive to many of the same factors that modulate biological peripersonal fields (Figures 2, 3). The structure of the artificial network subserving the agent’s actions is also reminiscent of its biological equivalents: it segregates into distinct sub-networks for appetitive versus avoidance actions (Figure 5, left panel). These arising properties indicate that the many different instances of biological peripersonal fields can be considered indexes of contact-action value, as originally hinted at by electrophysiology (115), and later suggested theoretically (16, 116, 117). Importantly, the formal explanation proposed here also makes multiple empirically testable predictions (see Box 3). This formalism has a further, theoretical implication: peripersonal fields can be combined into an egocentric map. The egocentric map provides a model of the world near the agent in terms of its future action values, which can be adapted to face novel tasks. This flexible model facilitates quick and efficient interactions with a constantly changing environment.

## Methods

### Model design

#### Base model

The agent’s environment was a grid world of 13×14 blocks (width [x] × height [y]), with recurring infinite boundary conditions on the left and right edges. In this world, an artificial neural network (ANN) controlled the x-position of its ‘limb’, located at y = 3. At each timestep, the ANN could either keep the limb still, or move it left or right. Simultaneously, an object moved in the world. When the object came into contact with the limb, the agent received a positive reward (+2).

The object spawned in at a random x-coordinate at the top of the world (i.e. at y = 14), and moved downward at a speed of 1 block per time step. In addition, in the same time step, the object had 50% chance to additionally move 1 block left, right, up, or down – a process simulating kinematic noise. After an object reached the bottom of the grid-world, a new object was spawned at the top of the grid-world. While staying still did not entail any reward (unless the object touched the limb), moving left or right had a cost of −0.001, to disincentivize random movement.

The ANN received world information through “proprioceptive” input reflecting the limb x-position, and “visual” input reflecting the object x and y positions. The network output was a value for each possible action (Q-value, see Box 1). Q-values were learned through Q-learning with experience replay (97). Each simulation lasted 4,000,000 time steps, each of which was stored as a state transition. At the end of the simulation, the network was further trained on stored state transitions to ensure near-optimal fitting of the value function, using 100 batches of 10,000 time steps each^10^.

#### Model variations

Most of the in-silico experiments described in the main text entail some variations to the base model environment described above. The exact variations are detailed in the description of each experiment, below. Here we only list the parameters that were varied.

1. <u>Number of limbs</u>. In some environments, we allowed the model to control an additional limb, placed one block below the base limb (i.e. at y = 2).
2. <u>Object velocity.</u> In some environments, objects with different velocity could spawn. Each time an object was spawned its y-velocity was set randomly between 1 and 3 (blocks/timestep). Similarly, its x-velocity was set randomly between −2 and +2.
3. <u>Input timesteps.</u> In the environments where the objects could have different velocities, the network was provided with a “memory” input: proprioceptive and visual information from the preceding timestep. This allowed the network to infer object velocity (118).
4. <u>Reward offered by objects</u>. In some environments, the reward consequent to contact with an object was set to either −2 or +4 instead of +2 of the base model. We note that we use the term ‘reward’ in the most general sense: thus, the agent can receive a negative reward, which can be understood as a punishment.
5. <u>Presence of a ‘tool’</u>. In some environments, we provided the agent with a ‘tool’, the effective part of which was located 4 blocks above the limb and measuring 1×1. The tool moved when the limb moved. When an object came into contact with the tool tip (i.e. the effective part of the tool), the agent experienced reward as if its limb had made contact with the object.
6. <u>Number of objects in the environment</u>. In some environments, two objects were simultaneously present. In such environments, we set the rewards to +2 for one object and −2 for the other, thus making the objects either ‘goals’ or ‘threats’, respectively.

We ran each of the experiments described below three times, each time training a separate neural network with different architectures. This procedure ensured that the results observed were not due to specific network architecture.

#### Combo Model

We also trained one agent in a model environment that included all variations described above. Where appropriate, we additionally ran the statistical tests on this ‘combo model’. This ensured that the results obtained in the simpler model variations also held for an agent that could display a wider range of environments with more complex behaviour.

All analyses were performed in Matlab. All codes used will be made available upon publication.

### Model #1 – Limb-centric response fields

#### Model design

In Model #1 we tested whether the action values (i.e. the ANN output), as well as the activity of single artificial units correlated with limb-object proximity. We ran this model 6 times, using a different network architecture each time. In 3 of the 6 instances the agent controlled 1 limb, and in the other 3 instances the agent controlled 2 limbs. When the agent controlled 2 limbs, the second limb was placed one row further down from the first limb (i.e. at y = 2). As for the first limb, the second limb could also be kept still or moved left or right at every timestep. Only goals (i.e. objects offering a +2 reward) were present in these environments.

To determine the number of network layers necessary for the agents to effectively interact with this environment, we assessed the performance of 12 networks composed of 1-12 layers. Each of these networks had 9 units per layer. We found that ANN composed of 4 and 5 layers were sufficient for near-optimal performance (~0.09 rewards per action) with one and two limbs, respectively. Network sizes were necessarily larger when controlling two limbs because the task was more complex; any of the three possible movements could be performed by each limb, thus requiring the agent to choose from a total of 9 actions (3×3) at every timepoint. As a result, we used network sizes of [12 10 8 6], [9 9 9 9] and [6 8 10 12] when controlling one limb, and [13 11 9 7 5], [9 9 9 9 9] and [5 7 9 11 13] when controlling two limbs. Each entry in these vectors represents the number of units in a layer, from the first (input) to the last (output) layer. The network shapes were chosen to demonstrate consistency of results across different network types, given that some of the analyses involved investigated the activity of individual units in the networks.

#### Statistical testing

To determine whether the Q-values correlated with the limb-object proximity, we calculated (separately for each model and limb) the Pearson correlation coefficient between the Q-values of each action and multiple measures of distance across all possible limb- and object-positions. These multiple measures were: absolute Euclidian limb-object distance, absolute limb-goal column-distance (i.e. x-axis distance), and absolute limb-goal row distance (i.e. y-axis distance). We considered a Q-value to be body-part centric if the FDR-corrected p-value was < 0.05 for all correlations across action types, number of models, and number of limbs. Q-values correlating with limb-object proximity constituted a peripersonal receptive field.

We performed identical analyses for each artificial unit to determine whether its activity also correlated with limb-object proximity (i.e. whether the neural activity was body-part centric), and could be therefore classified as peripersonal.

All analyses were performed separately for each of the six networks described above, as well as on another network trained on the ‘combo’ model (i.e. the model containing all the variations to the environment described in the section above). For the sake of conciseness, in the main text we report the highest, and thus weakest, p-values out of all Pearson’s correlation tests across all networks and actions.

To assess whether units with body-part centric receptive fields were homogeneously distributed across the ANNs layers, we calculated their proportion relative to the total number of units in each layer, separately for each ANN.

We finally ran a Linear Mixed Effect analysis (LME) to predict the proportion of peripersonal units, with relative layer depth as a fixed effect and network number as a random effect (as we assessed 7 separate networks).

### Model #2 – Kinematic effects

#### Model design

To test whether the peripersonal fields (1) expand when stimuli move faster, and (2) expand in the direction of incoming stimuli, we ran an additional model. The agent had a single limb, and was only exposed to goals. As detailed above (Model variation #2), every time a new goal was spawned, it was assigned a horizontal velocity. These velocities were [1 2 3] blocks per timestep horizontally and [−2 1 0 1 2] vertically. Thus, goals always moved downward, but with varying speeds [1 2 3] and stimulus directions [−2 1 0 1 2]. In addition, to provide the agent with information about the objects’ movement, the object position at the previous timestep was additionally provided to the network. We ran this model 3 times, using a different network architecture each time.

To determine the number of network layers necessary for the agents to satisfactorily interact with this environment, we assessed the performance of 12 networks composed of 1-12 layers. Each of these networks had 12 units per layer. We found that the ANN composed of 9 layers was sufficient for near-optimal performance (~0.1 rewards per action). Network sizes were necessarily larger than in Model #1 because the task was more complex: not only stimulus position but also speed and direction had to be taken into account. As a result, we used network sizes of [8 9 10 11 12 13 14 15 16], [12 12 12 12 12 12 12 12 12] and [16 15 14 13 12 11 10 9 8] units, from the first (input) to the last (output) layer. We chose these 3 network shapes to demonstrate consistency of results across different network types, given that part of the analysis involved the activity of individual units in the networks.

#### Statistical testing

To determine whether Q-value fields expanded as a function of the *velocity* with which objects fell downwards, we first averaged the Q-values across horizontal object speed ([−2 −1 0 1 2]) and possible actions. We then found the smallest vertical limb-object distance at which Q-values were above a particular threshold (the 90^th^ centile of Q-values in that condition). We calculated this distance for each of the three falling velocities [1 2 3], each limb position, each possible action, and each model. Next, we calculated the Pearson correlation coefficient ρ between this distance and the vertical object velocity. We considered a Q-field to be (1) *affected* by object velocity if the FDR corrected p-value was < 0.05 for all correlations, and (2) *expanded* by increasing object velocity if, additionally, the ρ value was positive (Figure 4).

To statistically test whether Q-value fields were affected by stimulus *direction*, we averaged Q-values across vertical object velocities [1 2 3], possible actions and vertical limb-object distances. We then calculated the asymmetry of the Q-fields as the horizontal (i.e. x-axis) distance between a) the center of mass of the Q-values distribution and b) the limb position. This distance was used as a proxy for lateral expansion of the Q-fields. We finally calculated the Pearson correlation coefficient between this lateral field expansion and the incoming stimulus direction, for each action (stay, move left, move right), and each of the 3 network architectures. We considered a Q-field to be affected by stimulus direction if the FDR corrected p-value was < 0.05 for all correlations.

All analyses were performed separately for each of the three network architectures described above, and also on another network trained on the ‘combo’ model (i.e. the model containing all the variations to the environment, as described in the ‘combo model’ section). For the sake of conciseness, in the main text we report the highest, and thus weakest, p-values out of all Pearson’s correlation tests across all networks and actions.

We performed identical analyses for each artificial unit to determine whether its neural activity could be described as a field which expands as a function of object velocity and direction, with only one difference: because artificial neural activity can be both positive and negative, this activity was (1) multiplied by the sign of its correlation with distance, and (2) shifted along the y-axis (i.e. baseline corrected by its lowest activity value) to be always positive. This manipulation ensured that the shape of the neural fields remain identical, while allowing the statistical analysis described above for Q-values. FDR correction was performed across units, layers and network architectures.

### Model #3 – Tool use, plastic and dynamic changes

#### Model design

To demonstrate the effects of tool use, we ran a model entailing only goals and a single limb. This time, however, we added the possibility for that the agent to hold a tool. This was modelled by an additional binary input to the network indicating whether the tool was present or not. We used two different types of tool: one effective and one ineffective. The ineffective tool had no effects on the environment or rewards. The effective tool, in contrast, resulted in a reward to the agent if one world block located 4 rows above the hand came into contact with a goal. This block represents the tip of the tool, effectively expanding the agent’s reach. Every time a new goal spawned, the agent had a 50% probability to ‘hold the tool’ until a new goal was spawned.

To demonstrate the reshaping of peripersonal fields due to tool use, we trained the agent and assessed its performance in two steps. First, we trained the agent with an ineffective tool (i.e. a condition in which contact between the tool tip and a goal did not offer any rewards). We then retrained the same agent with an effective tool.

To determine whether field reshaping occurred, we characterised the response fields of the agent before and after effective tool training, while they were and were not holding a tool. Specifically, we first averaged Q-values along the x-axis, and then calculated the number of Q-value peaks along the y-axis, for every possible limb position (a peak was defined as any block with a magnitude higher than its two neighbouring blocks along the y axis). Similarly, we also calculated the number of peaks for each artificial unit’s response fields.

We ran this model 3 times, using a different network architecture each time. The numbers of units and layers in these networks were determined as described in Model #2 section, but with a 0.5 probability of holding the tool and without multiple object speeds nor directions. As a result, we used network sizes of [8 10 12 14 16], [12 12 12 12 12] and [16 14 12 10 8] units, from the first to the last layer.

#### Statistical testing

To determine whether the Q-value fields had different shapes under any of the conditions, we performed a 2×2 ANOVA on the number of peaks in the Q-fields, with factors ‘tool training type’ (two levels: effective, innefective) and ‘tool presence’ (two levels: yes, no). To determine whether the Q-fields reshaped by holding an effective tool, we performed paired post-hoc Wilcoxon signed rank tests on the number of peaks, across limb positions and models.

We also calculated the proportion of ‘tool-use sensitive’ peripersonal units across neural network layers. We subtracted the number of peaks after training while holding the effective tool from the number of peaks before training, and labelled a unit as ‘tool-use-sensitive’ if this number was larger than 0 (i.e. if training resulted in a new peak in the response field). We then calculated the fraction of such units for each layer and neural network separately. We finally ran a Linear Mixed Effect analysis (LME) to predict the proportion of tool-sensitive units, with layer depth as a fixed effect and network number as a random effect.

### Model #4 – Valence effects

#### Model designs

##### Expansion of receptive fields (4a)

To test the effects of valence magnitude, we ran a model entailing two different types of goals, and a single limb. The rewards for making contact with the first and second goal types were +2 and +4, respectively. The two types of goals moved independently. To probe the effects of valence on movement choice more effectively, the cost of moving (−0.1) was higher than in the other models (−0.001). The agent received “visual” input reflecting the x and y position of each goal type separately. The numbers of units and layers in these networks were determined as described in Model #2. As a result, we used network sizes of [6 10 14 18], [12 12 12 12] and [18 14 10 6] units, from the first to the last layer.

##### Separation of movement classes (4b)

To test the effect of negative valence stimuli, we used a model identical to model 5a, but entailing both goals and threats. The rewards for contacting these stimuli were set to +2 and – 2 respectively. Network sizes were also as above ([6 10 14 18], [12 12 12 12] and [18 14 10 6] units, from first to last layer). Because we found that these each of these three networks split into two distinguishable sub-networks (one responding more to threats, and one to goals; see Results), we assessed the reliability of the generation of such sub-networks. To do so, we trained from scratch three new networks of the same sizes on new data, 15 times per network size (so training 45 networks in total). To ensure that the results were not due to the specific minimal network sizes, we then trained three additional new networks of larger sizes ([6 10 12 14 16 18], [12 12 12 12 12 12 12] and [18 16 14 12 10 8 6]), again 15 times per network size. Results reported in the main text refer to this final training of larger networks, because (1) statistical results using more units are more robust, and (2) this last analysis represents the final validation of the results. The results from the smaller networks, reported below in the ‘statistical testing’ section, were qualitatively similar.

#### Statistical testing

##### Expansion of receptive fields (4a)

First, to test whether Q-value magnitudes are larger in response to stimuli with higher valence, we performed sign rank tests between the Q-values obtained with low-reward and high-reward goals. We performed these tests for each action (move left, move right and stay), and each of the 3 network architectures. We considered a Q-field to be affected by reward magnitude if the FDR-corrected p-value was < 0.05 for all comparisons.

Second, to test whether actions related to a high-reward object are performed when the object is at greater distances from the body as compared to a low-reward object, we shifted Q-values by the minimum value, to make them all positive. We then compared the Q-values in the y-direction to a normally-distributed noisy threshold for action initiation: the threshold was centric at the 90^th^ centile of Q-values, with a standard deviation of 25% of the 90^th^ centile. The proportion of initiated contact-related actions was calculated by entering the Q-value into the normal cumulative distribution function of the threshold distribution. We performed this analysis in all models, for all limb positions and actions. Next, we compared the distance at which the proportion of contact actions was above 0.5 between stimuli of the two reward magnitudes, using a Wilcoxon signed-rank test for each network architecture. A difference on this test indicates that higher-reward stimuli result in actions at further distances from the body.

Third, we assessed the effects of reward magnitude on artificial unit response. For each unit and network, we calculated the variability in neural activity for all object and limb positions. We did this separately for high- and low-reward objects. We then assessed whether units were overall more responsive to higher-reward objects by calculating the difference in variability between high- and low-valence objects, dividing this difference by the sum of the variance across reward magnitudes, and comparing the distribution of these normalized differences against zero using a Wilcoxon signed rank test.

##### Separation of movement classes (4b)

First, we verified that both Q-values and neural activity correlated with limb-object proximity. We performed the same analyses described for Model #1, but this time also on Q-values and neural activities resulting from negative reward objects (i.e. threats). To perform the analyses relative to goals, we averaged the Q-values across all possible threat positions, and vice-versa.

Next, we investigated the differences in agents’ behaviour in response to goals and threats. We labelled the agents’ behaviour as follows: −1 if the limb moved to intercept the object (i.e. if it moved to reduce the x-axis distance to the goal or the threat), 0 if the limb did not move, and +1 if the limb moved to avoid the object (i.e. if it moved to increase the x-axis distance to the goal or the threat), for every network architecture, object position, and limb position. We then performed a Wilcoxon signed rank test on these movement types, comparing the presence of goals with the presence of threats, across models and object positions.

Subsequently, we used two different approaches to test whether distinguishable sub-networks arose in these agents.

First, we assessed whether units of a given type were more strongly connected to other units of the same type. To do so, we (1) created ***c***, a measure of how much each unit *i* in a network prefers either positive or negative objects. This measure of preference ***c*** was defined as the difference in absolute covariance between a unit’s activity ***a***_*i*_ and the distance ***d***_*i*_ between the limb and either types of object:

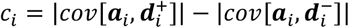

Where 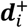 and 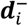 are vectors of distances between the limb and either positive-reward or negative-reward objects, respectively. We then (2) assessed whether each unit is more connected to other units with similar preference ***c***: we calculated the Pearson correlation coefficient between all ***c***, and the weighted preferences of all units connected to them, *W****c***. The weights *W*_*ij*_ were the input weights from each unit *i* to each unit *j*, z-scored across all inputs for each *j*. We considered a network to contain distinguishable sub-networks (according to this first metric) if the FDR-corrected correlation p-value was < 0.05.

A second approach to test whether distinguishable sub-networks arose, we quantified the *spatial* sub-network structure, assuming that more strongly connected units are more likely to be closer to each another. To do so, we projected each network onto a 2D plane as a weighted graph using the matlab *digraph* function, with edge length set to the inverse of the weight between every pair of units. Next, we assigned the neural preference *c*_*i*_ to the position of each unit *i*, and applied a 2D gaussian filter (*σ* = [5, 5]) across the graph. In this way, areas of the graph with large groups of similarly-classified units would have high absolute values. Finally, we summed these absolute values (i.e. the values of filtered similarity index) over the entire graph. We then permuted the location of the *c*_*i*_ values 100 times, filtered and summed them, and finally compared the original sum against this null distribution. We considered the networks to be more structured than by chance if the difference between the real and permuted values was > 0 (assessed using an un-paired t-test).

All analyses were performed on the 45 (15×3) networks described above, as well as on the ‘combo’ network trained on the more realistic environment. As for all previous models, in the main text we report the highest, and thus weakest, p-values across all networks.

##### Small-network results

Both goals and threats resulted in action-value fields graded with limb-object proximity and anchored to limb position (p ≤8.98 ×10^−4^, ρ ≤0.124 for positive valence stimuli; p ≤5.59 ×10^−6^, ρ ≥0.163 for negative valence stimuli; FDR-corrected Pearson correlation tests; Figure 5). Behaviourally, the agents moved their limb away from threats and towards goals (p ≤5.82 ×10^−4^, |Z| ≥3.44). The activity of a substantial proportion of units within the network correlated with the proximity of at least one stimulus type (79 ±7%). In addition, units showed preference for a given stimulus types. 60 ±12% of neuronal activities correlated with goal proximity, 55 ±12% with threat proximity, and 36 ±10% with both. In almost every network architecture, two sub-networks emerged: goal preferring units were more strongly connected to each other and vice-versa (p ≤0.05 in 28 out of 48 tested networks, FDR-corrected Pearson correlation). Importantly, there was more sub-network structure than expected by chance across all networks (p ≤5.59 ×10^−6^, T =7.29; un-paired t-test).

### Model #5 – Transferable Successor Features

#### Model design

To demonstrate that Q-fields can be used as a set of successor features for near-body interactions, we used two different environments. In one environment the agent was only exposed to threats (yielding a −2 reward upon contact). In the other the agent was only exposed to goals (yielding +2 reward upon contact). To determine network sizes we used the same approach as for Model #1. As a result, we used network sizes of [12 10 8 6], [9 9 9 9] and [6 8 10 12] units, from the first to the last layer. We used these same networks to demonstrate that successor features encode impact probability (see *Reconstruction of hit probability*, below). To make the task of reconstructing a negative-valence value field harder, we altered the dynamics of the positive-reward objects: instead of a 50% chance of moving an additional block up, down, left or right, goals had a 33% chance of moving an additional block left or right, and never moved an additional block up or down.

#### Statistical testing

##### Transferable successor features

We reconstructed Q-values in response to *negative*-reward stimuli (threats) from Q-values in response to *positive*-reward objects (goals). To do so, we first retrieved all Q-values calculated by an agent that had only been exposed to goals, for all stimulus and limb positions. We did the same for all Q-values calculated by an agent that had only been exposed to threats. We then linearly reconstructed the latter (i.e. the Q-value to *negative*-reward stimuli for ‘stay’ action) using the former (i.e. the Q-values to *positive*-reward stimuli for ‘stay’, ‘move left’, and ‘move right’ actions).

We subsequently calculated the strength of the relationship between the reconstructed threat Q-values and the original threat Q-values using a Pearson correlation. To demonstrate that body-part centric *neural* activity (i.e. value function *approximations*) can be exploited in the same manner, we repeated this analysis a second time. The only difference was that instead of using goals Q-values to reconstruct threats Q-values, we took the neural activities from the later layers of the neural network that calculated Q-values in response to goals (Figure 5).

We performed these comparisons between the two models of each network architecture, but trained on either goals or threats. We considered the reconstructed Q-fields to be good approximations of the original values if the FDR corrected p-value was < 0.05 for all tests.

##### Reconstruction of hit probability

We also reconstructed the probability that any object would make an impact with the limb (i.e. the probability of experiencing touch given vision, 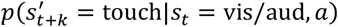 from existing Q-values. We first calculated the actual impact probability of an object using the observations stored from the agent training (note that objects disappeared after they either contacted the limb or touched the bottom of the world [y=0], and so impact probability was always ≤1.) We subsequently linearly reconstructed the hit probabilities for all world states using the goal Q-values, and calculated the correlation strength as we did it for reconstructed threat Q-values. Again, we performed the same analysis using the neural activity from the later layers of the network. We performed these reconstructions using all architectures of the models trained with goals. We considered the reconstructed hit probabilities to be good approximations of the original values if the FDR-corrected p-value was < 0.05.

### Model #6 – Psychophysical responses

#### Model design

To demonstrate that behavioural responses that do not necessarily entail avoiding or creating contact with objects can nonetheless be body-part centric, we trained a new ANN to perform a different task. This network controlled one limb as described above, but contact with environmental objects was not rewarded. Instead, the agent was rewarded for performing a correct ‘button press’ action in response to an occasional extra ‘tactile’ input given to the ANN concomitantly to the standard visual and proprioceptive input. Crucially, the tactile input was noisy, i.e. it fluctuated randomly around 1 and 0 (when tactile input was present and absent, respectively).

The agent was rewarded when it responded correctly to the tactile input with a button press. If the agent responded when there was no tactile input, it was punished. We calculated the average response rate to these noisy tactile stimuli. This model, besides simulating a tactile detection task, can also be interpreted as simulating a reaction time task: assuming that better detection leads to faster reaction times, a high average response rate indicates a short reaction time and vice versa.

The network performing the reaction time task had a crucial quirk: it contained a pre-trained subnetwork (specifically, the network described in Model #1) that returned the Q-values of moving under the assumption that contact with visual stimuli would result in a positive reward, even if in the current task contact was irrelevant to receive a reward (Figure 3).

To simulate the results of rTMS lesion experiments in humans (59), we calculated the average response rate under two conditions: 1) leaving the network intact, as described above, and 2) lesioning the network by setting the output of the pre-trained module to zero for any input condition, thus mimicking an rTMS lesion of the PPC in humans.

#### Statistical testing

We tested whether the network lesioning removed the proximity effect on reaction times. For the sake of analogy to how real rTMS lesion data are presented (Figure 3), we grouped the reaction times according to whether the visual stimulus was ‘Near’ or ‘Far’ from the limb (59). Near and Far responses were the average reaction times when stimuli were 1-5 and 7-11 blocks away from the limb along the y-axis, respectively. We calculated these measures for each of the 13 possible limb position. Next, for each limb position, we subtracted the Far from the Near responses, to obtain a measure of the shortening of reaction times as a function of stimulus proximity. We compared this reaction time shortening to zero using a Wilcoxon signed rank test, as well as between the intact and rTMS lesioned neural networks using a Wilcoxon rank sum test.

## Supplementary Materials

### Supplementary Figure

**Figure S1.**
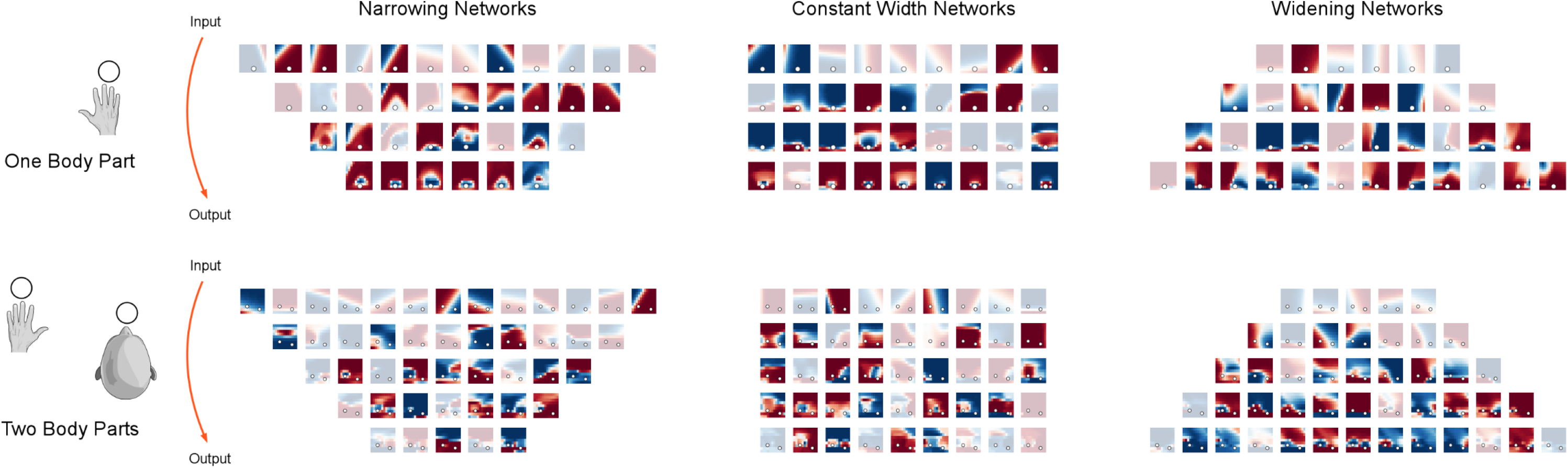
Individual neural response fields in all architectures used for model 1. Each heatmap show the activity of a network unit as a function of spatial position relative to the body part. Full-colour heatmaps came from units classed as peripersonal, while grayed-out heatmaps did not. The proportion of peripersonal units per layer increased as a function of layer depth

### Supplementary Videos

Supplementary videos are available at https://doi.org/10.6084/m9.figshare.20449758.v3

#### Video legends

**Supplementary Video SV1.**
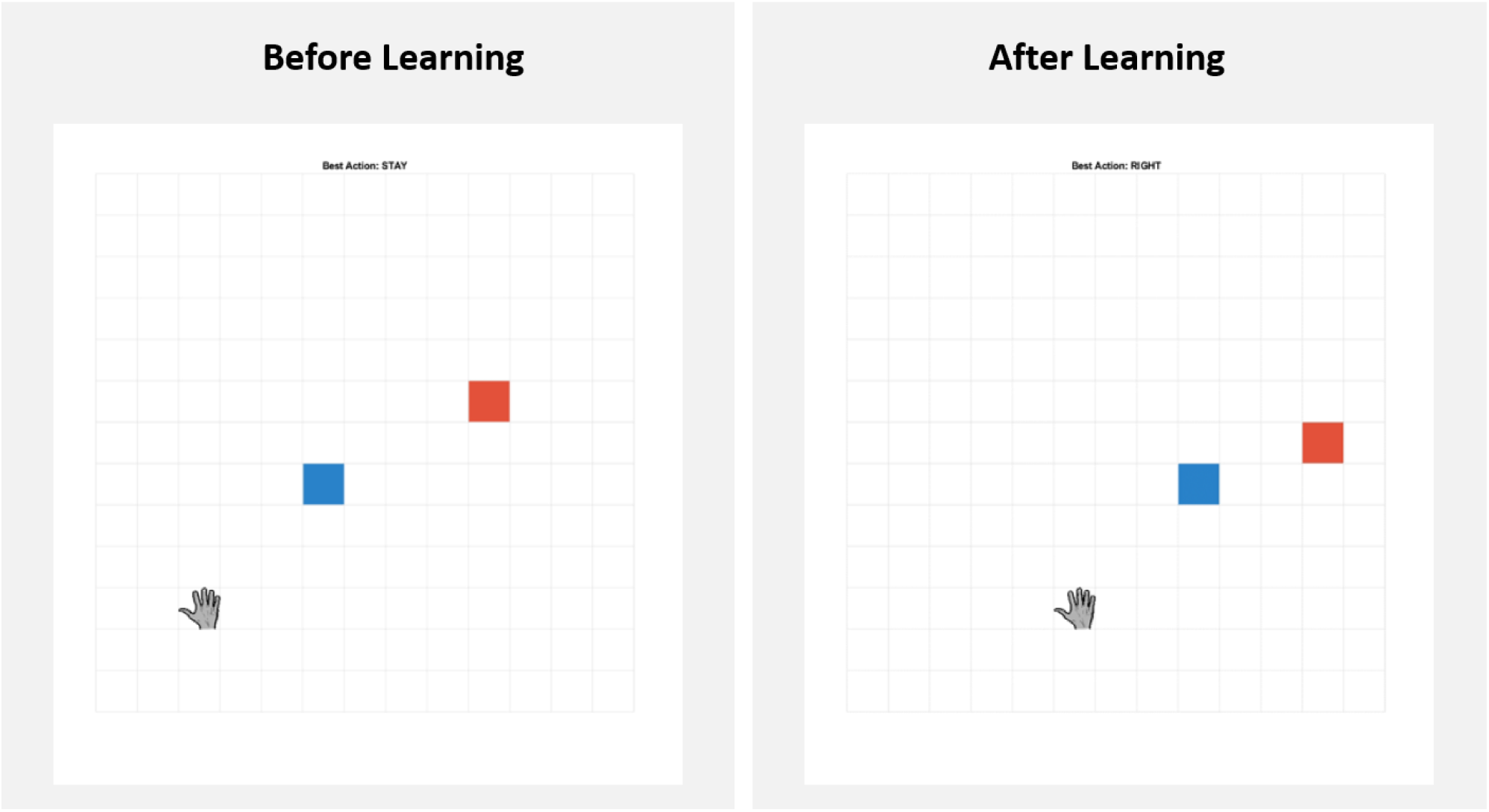
Learning agent. Agents are trained to move their limbs (grey hand) to avoid threats (red squares) and intercept goals (blue squares). Video shows instances of an example agent being trained in an environment where threats move less predictably than goals. Before training, the agent’s behaviour is random (left panel). After some training, the agent intercepts goals, but only when it is unlikely to be hit by a threat.

**Supplementary Video SV2.**
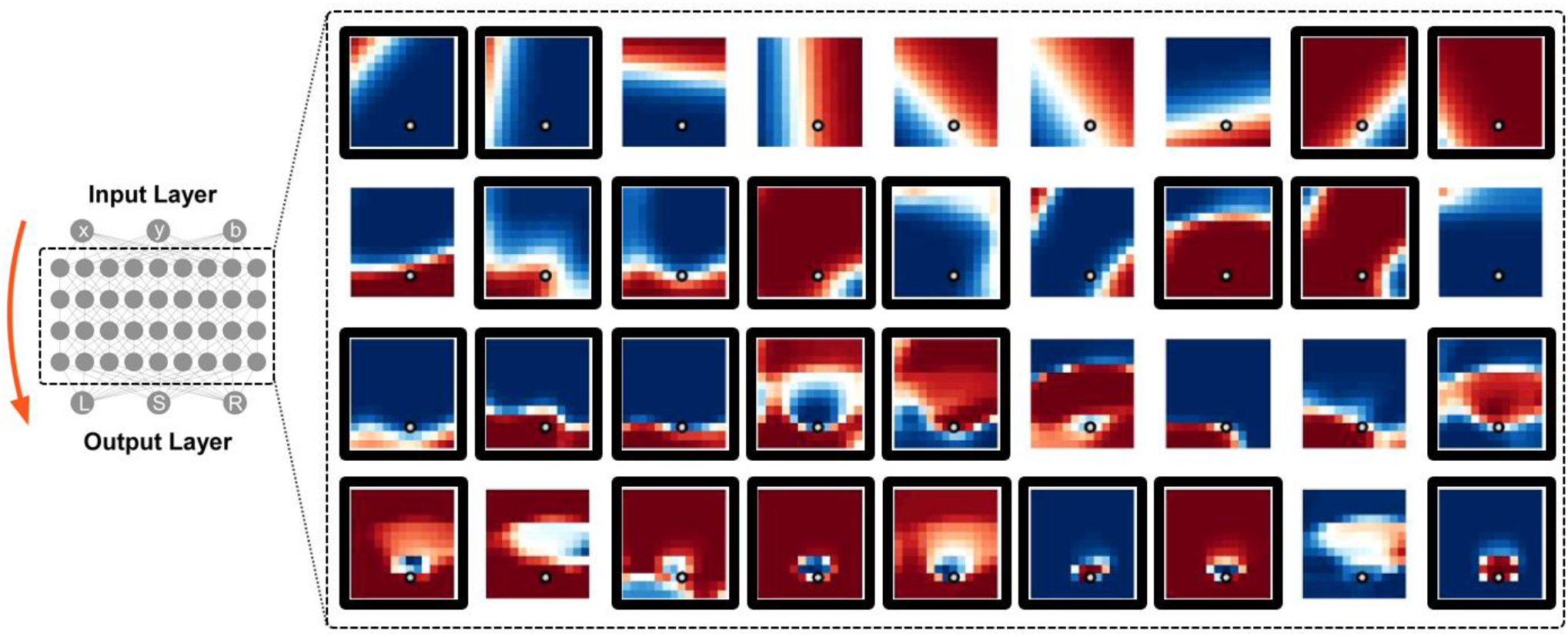
Shifting fields. A large portion of artificial neurons display body-part centric responses that shift with the location of the limb. Receptive fields of each artificial neuron are shown as colour maps. ‘Peripersonal’ receptive fields are highlighted by a black box.

## Footnotes

1 The responses of interest here are egocentric in terms of *proximity*, and thus *distance*. In this article, we do not discuss the many neural responses which are egocentric in terms of *angle* only (119–123). We therefore use the term ‘peripersonal’ to refer to a response field that is egocentric but *also* centred around and anchored to a body-part.

2 These behavioural responses include cross-modal extinction in brain-damaged patients (31), crossmodal congruency distraction effects (124), performance in line bisection tasks (33), reaction times to tactile stimuli during simultaneous visual or auditory stimulation (34, 35, 113), temporal order judgements (36), reachability and semantic estimates (37, 38), spatial demonstratives (39), defensive reflexes (40, 125), and TMS-evoked motor potentials (59).

3 Here we use the term ‘reward’ in its reinforcement learning sense, allowing it to have both positive (i.e. a prize) and negative meaning (i.e. a punishment) depending on its sign.

4 You might, however, still feel the urge to walk away from the wasp. Thus, when the wasp is at an intermediate distance, the value of walking away is substantially higher than the value of moving one limb. This illustrates the important point that there are different peripersonal fields for different actions: the ‘walk away field’ would presumably be centred on the whole body and expand far into space, while the ‘move hand out the way’ field would be centred on the hand and only expands is less expansive (16).

5 *Plastic* remapping here refers to the change of a neural response field after extended tool use, while *dynamic* remapping refers to the immediate change as soon as the tool is picked up (86)

6 We are purposefully modelling tool use in a highly abstracted manner: here, a tool is simply something the agent can use to modify the effect of its actions on the environment. Specifically, if the agent ‘holds a tool’, the agent receives a reward when objects are in contact with either its limb, as usual, or the location of the tool tip.

7 Unfortunately, studies on single-cell recordings never statistically tested the effects of valence, but their authors did recommend using stimuli with an inherent interest to the monkey – such as food, or novel objects (77, 126).

8 This holds true because (i) at a given timepoint, the value of any action assumes that the agent will perform the optimal actions at all future time points (see Box 1), and (ii) the optimal actions are different when facing a threat or a goal.

9 Although very recently, an impact prediction model has touched on the importance of motor repertoire: that model suggests that impact prediction better fits peripersonal data when one takes into account the time it takes to perform an action with the limb in question (104).

10 The scope of this article was not to create an algorithm that optimally learns particular Q-values. Therefore, the parameters used to learn Q-values were not rigorously optimized with respect to computational speed or performance. These parameters were chosen because they gave satisfactory performance, without substantial time investment.

## References

1. E. I. Moser, E. Kropff, M.-B. Moser, Place Cells, Grid Cells, and the Brain’s Spatial Representation System. Annual Review of Neuroscience 31, 69–89 (2008).

2. O’Keefe, J. Dostrovsky, The hippocampus as a spatial map. Preliminary evidence from unit activity in the freely-moving rat. Brain Res 34, 171–175 (1971).

3. P. J. Best, A. M. White, A. Minai, Spatial processing in the brain: the activity of hippocampal place cells. Annual Review of Neuroscience 24, 459–86 (2001).

4. A. Arleo, W. Gerstner, Spatial cognition and neuro-mimetic navigation : a model of hippocampal place cell activity. Biol Cybern 83, 287–299 (2000).

5. A. Banino, et al., Vector-based navigation using grid-like representations in artificial agents. Nature 557, 429–433 (2018).

6. C. Barry, N. Burgess, Learning in a geometric model of place cell firing. Hippocampus 17, 786–800 (2007).

7. S. N. Weber, H. Sprekeler, Learning place cells, grid cells and invariances with excitatory and inhibitory plasticity. Elife 7, 1–41 (2018).

8. K. L. Stachenfeld, M. M. Botvinick, S. J. Gershman, Supplemental Information for Design Principles of the Hippocampal Cognitive Map. Advances in Neural Information Processing Systems 27 1, 1–9 (2014).

9. K. L. Stachenfeld, M. M. Botvinick, S. J. Gershman, The hippocampus as a predictive map. Nature Neuroscience 20, 1643–1653 (2017).

10. B. E. Pfeiffer, D. J. Foster, Hippocampal place-cell sequences depict future paths to remembered goals. Nature 76(2013).

11. I. Momennejad, et al., The successor representation in human reinforcement learning. Nature Human Behaviour 1, 680–692 (2017).

12. E. M. Russek, I. Momennejad, M. M. Botvinivk, S. J. Gershman, N. D. Daw, Predictive representations can link model-based reinforcement learning to model-free mechanisms. Momennejad, Ida Botvinick, Matthew M. Gershman, Samuel J. Daw, Nathaniel D.PLoS Computational Biology 13(2017).

13. M. M. Garvert, R. J. Dolan, T. E. J. Behrens, A map of abstract relational knowledge in the human hippocampal–entorhinal cortex. Elife 6, 1–20 (2017).

14. D. Aronov, R. Nevers, D. W. Tank, Mapping of a non-spatial dimension by the hippocampal–entorhinal circuit. Nature Publishing Group 543, 719–722 (2017).

15. R. A. Epstein, E. Z. Patai, J. B. Julian, H. J. Spiers, The cognitive map in humans : spatial navigation and beyond (2017) https://doi.org/10.1038/nn.4656.

16. R. J. Bufacchi, G. D. Iannetti, An Action Field Theory of Peripersonal Space. Trends in Cognitive Sciences 22, 1076–1090 (2018).

17. M. S. Graziano, D. F. Cooke, Parieto-frontal interactions, personal space, and defensive behavior. Neuropsychologia 44, 845–859 (2006).

18. E. Ladavas, Functional and dynamic properties of visual peripersonal space. Trends Cogn Sci 6, 17–22 (2002).

19. J.-P. Noel, O. Blanke, A. Serino, From multisensory integration in peripersonal space to bodily self-consciousness: from statistical regularities to statistical inference. Ann N Y Acad Sci 1426, 146–165 (2018).

20. N. P. Holmes, C. Spence, The body schema and the multisensory representation(s) of peripersonal space. Cogn Process 5, 94–105 (2004).

21. C. L. Colby, J. R. Duhamel, M. E. Goldberg, Ventral intraparietal area of the macaque: anatomic location and visual response properties. J Neurophysiol 69, 902–914 (1993).

22. J. R. Duhamel, C. L. Colby, M. E. Goldberg, Ventral intraparietal area of the macaque: congruent visual and somatic response properties. J Neurophysiol 79, 126–136 (1998).

23. M. S. Graziano, C. G. Gross, A bimodal map of space: somatosensory receptive fields in the macaque putamen with corresponding visual receptive fields. Experimental brain research. Experimentelle Hirnforschung. Experimentation cerebrale 97, 96–109 (1993).

24. G. Rizzolatti, C. Scandolara, M. Matelli, M. Gentilucci, Afferent properties of periarcuate neurons in macaque monkeys. II. Visual responses. Behavioural Brain Research 2, 147–163 (1981).

25. J. Hyvarinen, A. Poranen, Function of the parietal associative area 7 as revealed from cellular discharges in alert monkeys. Brain 97, 673–692 (1974).

26. F. Bremmer, et al., Polymodal motion processing in posterior parietal and premotor cortex. Neuron 29, 287–296 (2001).

27. T. R. Makin, N. P. Holmes, E. Zohary, Is that near my hand? Multisensory representation of peripersonal space in human intraparietal sulcus. J Neurosci 27, 731–40 (2007).

28. J. Cléry, O. Guipponi, S. Odouard, C. Wardak, S. Ben Hamed, Cortical networks for encoding near and far space in the non-human primate. Neuroimage 176, 164–178 (2018).

29. Y. Wamain, F. Gabrielli, Y. Coello, EEG μ rhythm in virtual reality reveals that motor coding of visual objects in peripersonal space is task dependent. Cortex 74, 20–30 (2016).

30. C. F. Sambo, B. Forster, An ERP investigation on visuotactile interactions in peripersonal and extrapersonal space: evidence for the spatial rule. J Cogn Neurosci 21, 1550–9 (2009).

31. A. Farnè, E. Làdavas, Auditory peripersonal space in humans. J Cogn Neurosci 14, 1030–1043 (2002).

32. C. Spence, F. Pavani, A. Maravita, N. Holmes, Multisensory contributions to the 3-D representation of visuotactile peripersonal space in humans: Evidence from the crossmodal congruency task. Journal of Physiology Paris 98, 171–189 (2004).

33. M. E. McCourt, M. Garlinghouse, Asymmetries of visuospatial attention are modulated by viewing distance and visual field elevation: pseudoneglect in peripersonal and extrapersonal space. Cortex 36, 715–731 (2000).

34. A. M. de Haan, M. Smit, S. Van der Stigchel, H. C. Dijkerman, Approaching threat modulates visuotactile interactions in peripersonal space. Experimental Brain Research 234, 1875–1884 (2016).

35. E. Canzoneri, E. Magosso, A. Serino, Dynamic sounds capture the boundaries of peripersonal space representation in humans. PLoS One 7, e44306 (2012).

36. A. L. De Paepe, G. Crombez, C. Spence, V. Legrain, Mapping nociceptive stimuli in a peripersonal frame of reference: evidence from a temporal order judgment task. Neuropsychologia 56, 219–228 (2014).

37. J. Bourgeois, Y. Coello, Effect of visuomotor calibration and uncertainty on the perception of peripersonal space. Attention, Perception, and Psychophysics 74, 1268–1283 (2012).

38. S. Kalénine, Y. Wamain, J. Decroix, Y. Coello, Conflict between object structural and functional affordances in peripersonal space. Cognition 155, 1–7 (2016).

39. M. Caldano, K. R. Coventry, Spatial demonstratives and perceptual space: To reach or not to reach? Cognition 191(2019).

40. V. Versace, S. Campostrini, L. Sebastianelli, L. Saltuari, M. Kofler, Modulation of exteroceptive electromyographic responses in defensive peripersonal space. Journal of Neurophysiology 121, 1111–1124 (2019).

41. B. A. Stettler, L. E. Thomas, Visual processing is biased in peripersonal foot space. Attention, Perception, & Psychophysics, 298–305 (2017).

42. F. de Vignemont, A. Serino, H. Y. Wong, A. Farne, The world at out fingertips. A multidisciplinary exploration of peripersonal space, ! (Oxford University Press, 2021) https://doi.org/10.1093/oso/9780198851738.001.0001.

43. E. Làdavas, A. Farnè, Visuo-tactile representation of near-the-body space. J Physiol Paris 98, 161–70 (2004).

44. A. Serino, Peripersonal space (PPS) as a multisensory interface between the individual and the environment, defining the space of the self. Neuroscience and Biobehavioral Reviews 99, 138–159 (2019).

45. J. Cléry, et al., The prediction of impact of a looming stimulus onto the body is subserved by multisensory integration mechanisms. The Journal of Neuroscience 37, 10656–10670 (2017).

46. T. Bertoni, E. Magosso, A. Serino, From statistical regularities in multisensory inputs to peripersonal space representation and body ownership: Insights from a neural network model. European Journal of Neuroscience 53, 611–636 (2021).

47. Z. Straka, M. Hoffmann, Learning a Peripersonal Space Representation as a Visuo-Tactile Prediction Task. 1, 101–109 (2017).

48. R. J. Bufacchi, M. Liang, L. D. Griffin, G. D. Iannetti, A geometric model of defensive peripersonal space. Journal of Neurophysiology 115, 218–225 (2016).

49. A. Roncone, M. Hoffmann, U. Pattacini, L. Fadiga, G. Metta, Peripersonal space and margin of safety around the body: Learning visuo-tactile associations in a humanoid robot with artificial skin. PLoS ONE 11(2016).

50. E. Magosso, M. Zavaglia, A. Serino, G. di Pellegrino, M. Ursino, Visuotactile representation of peripersonal space: a neural network study. Neural Comput 22, 190–243 (2010).

51. R. J. Bufacchi, G. D. Iannetti, “What do ‘peripersonal space measures’ really reflect? The action field perspective.” in The World at Our Fingertips: A Multidisciplinary Exploration of Peripersonal Space., (Oxford University Press, 2021), pp. 155–180.

52. A. Serino, Peripersonal space (PPS) as a multisensory interface between the individual and the environment, defining the space of the self. Neuroscience & Biobehavioral Reviews (2019) https://doi.org/10.1016/J.NEUBIOREV.2019.01.016.

53. G. Rizzolatti, L. Fadiga, L. Fogassi, V. Gallese, The space around us. Science 277, 190–191 (1997).

54. L. Fogassi, V. Gallese, “Action as a binding key to multisensory integration” in The Handbook of Multisensory Processes, (2004), pp. 425–441.

55. H. C. Dijkerman, A. Farnè, Sensorimotor and social aspects of peripersonal space. Neuropsychologia 70, 309–312 (2015).

56. S. B. Hunley, S. F. Lourenco, What is peripersonal space ? An examination of unresolved empirical issues and emerging findings. Cognitive Science 9(2018).

57. E. Blini, A. Farnè, C. Brozzoli, F. Hadj-Bouziane, “Close is better: Visual perception in peripersonal space.” in The World at Our Fingertips: A Multidisciplinary Exploration of Peripersonal Space., (Oxford University Press, 2021), pp. 47–60.

58. G. di Pellegrino, E. Làdavas, Peripersonal space in the brain. Neuropsychologia 66, 126–133 (2015).

59. A. Serino, E. Canzoneri, A. Avenanti, Fronto-parietal areas necessary for a multisensory representation of peripersonal space in humans: An rTMS study. J Cogn Neurosci 23, 2956–2967 (2011).

60. D. Lloyd, N. Roberts, Role for Human Posterior Parietal Cortex in Visual Processing of Aversive Objects in Peripersonal Space. J Neurophyiol 95, 205–214 (2006).

61. C. Brozzoli, G. Gentile, H. H. Ehrsson, That’s near my hand! Parietal and Premotor coding of hand-centered space contributes to localization and self-attribution of the hand. Journal of Neuroscience 32, 14573–14582 (2012).

62. E. J. Kim, et al., Alterations of Hippocampal Place Cells in Foraging Rats Facing a “Predatory” Threat. Current Biology, 1–6 (2015).

63. M. Berger, P. Neumann, A. Gail, Peri-hand space expands beyond reach in the context of walk-and-reach movements. Scientific Reports, 1–12 (2019).

64. J. P. Noel, et al., Full body action remapping of peripersonal space: The case of walking. Neuropsychologia 70, 375–384 (2014).

65. R. Somervail, et al., Movement of environmental threats modifies the relevance of the defensive eye-blink in a spatially-tuned manner. Scientific Reports 9(2019).

66. L. Fogassi, et al., Coding of peripersonal space in inferior premotor cortex (area F4). J Neurophysiol 76, 141–157 (1996).

67. M. K. Huijsmans, A. M. de Haan, B. C. N. Müller, H. C. Dijkerman, H. T. van Schie, Knowledge of Collision Modulates Defensive Multisensory Responses to Looming Insects in Arachnophobes. Journal of Experimental Psychology: Human Perception and Performance 48, 1–7 (2022).

68. A. Iriki, M. Tanaka, Y. Iwamura, Coding of modified body schema during tool use by macaque postcentral neurones. Neuroreport 7, 2325–2330 (1996).

69. A. Farnè, A. Iriki, E. Làdavas, Shaping multisensory action-space with tools: Evidence from patients with cross-modal extinction. Neuropsychologia 43, 238–248 (2005).

70. A. Rossetti, D. Romano, N. Bolognini, A. Maravita, Dynamic expansion of alert responses to incoming painful stimuli following tool use. Neuropsychologia 70, 486–494 (2015).

71. A. Berti, F. Frassinetti, When far becomes near: remapping of space by tool use. J Cogn Neurosci 12, 415–420 (2000).

72. N. P. Holmes, G. A. Calvert, C. Spence, Tool use changes multisensory interactions in seconds: Evidence from the crossmodal congruency task. Experimental Brain Research 183, 465–476 (2007).

73. N. P. Holmes, Does tool use extend peripersonal space? A review and re-analysis. Experimental Brain Research 218, 273–282 (2012).

74. A. Serino, M. Bassolino, A. Farnè, E. Làdavas, Extended multisensory space in blind cane users. Psychol Sci 18, 642–8 (2007).

75. A. Maravita, A. Iriki, Tools for the body (schema). Trends in Cognitive Sciences 8, 79–6 (2004).

76. M. A. Umilta, et al., When pliers become fingers in the monkey motor system. Proceedings of the National Academy of Sciences 105, 22902213 (2008).

77. M. Graziano, The spaces between us: A story of neuroscience, evolution, and human nature, 1st Ed. (Oxford University Press, 2017).

78. V. B. Mountcastle, J. C. Lynch, A. Georgopoulos, H. Sakata, C. Acuna, Posterior parietal association cortex of the monkey: command functions for operations within extrapersonal space. Journal of Neurophysiology 38, 871–908 (1975).

79. C. L. Colby, M. E. Goldberg, Space and attention in parietal cortex. Annual Review of Neuroscience 22, 319–349 (1999).

80. F. Ferri, A. Tajadura-Jiménez, A. Väljamäe, R. Vastano, M. Costantini, Emotion-inducing approaching sounds shape the boundaries of multisensory peripersonal space. Neuropsychologia 70, 468–475 (2015).

81. C. Spaccasassi, D. Romano, A. Maravita, Everything is worth when it is close to my body: How spatial proximity and stimulus valence affect visuo-tactile integration. Acta Psychologica 192, 42–51 (2019).

82. E. Vagnoni, S. F. Lourenco, M. R. Longo, Threat modulates perception of looming visual stimuli. Curr Biol 22, R826–7 (2012).

83. Y. Coello, F. Quesque, M. F. Gigliotti, L. Ott, J. L. Bruyelle, Idiosyncratic representation of peripersonal space depends on the success of one’s own motor actions, but also the successful actions of others! PLoS ONE 13, 1–30 (2018).

84. A. Zanini, R. Salemme, C. Brozzoli, Associative learning in Peripersonal Space : fear responses are acquired in hand-centered. Journal of Neurophysiology (2021).

85. F. de Vignemont, G. D. Iannetti, How many peripersonal spaces? Neuropsychologia 70, 327–334 (2015).

86. J. Cléry, O. Guipponi, C. Wardak, S. Ben Hamed, Neuronal bases of peripersonal and extrapersonal spaces, their plasticity and their dynamics: Knowns and unknowns. Neuropsychologia 70, 313–326 (2015).

87. C. Spaccasassi, C. H. Dijkerman, A. Maravita, O. Ferrante, M. C. De Jong, Body – Space Interactions : Same Spatial Encoding but Different Influence of Valence for Reaching and Defensive Purposes. 1–18 (2021).

88. D. F. Cooke, M. S. Graziano, Super-flinchers and nerves of steel: defensive movements altered by chemical manipulation of a cortical motor area. Neuron 43, 585–93 (2004).

89. M. S. A. Graziano, C. S. R. Taylor, T. Moore, Complex movements evoked by microstimulation of precentral cortex. Neuron 34, 841–851 (2002).

90. A. Zanini, et al., Patterns of multisensory facilitation distinguish peripersonal from reaching space. 1–36 (2020).

91. R. Caminiti, et al., Computational architecture of the parieto-frontal network underlying cognitive-motor control in monkeys. eNeuro 4, ENEURO.0306-16.2017 (2017).

92. T. N. Aflalo, M. S. A. Graziano, Possible Origins of the Complex Topographic Organization of Motor Cortex : Reduction of a Multidimensional Space onto a Two-Dimensional Array. 26, 6288–6297 (2006).

93. P. Cisek, J. F. Kalaska, Neural mechanisms for interacting with a world full of action choices. Annual Review of Neuroscience 33, 269–298 (2010).

94. M. Graziano, The Intelligent Movement Machine: An Ethological Perspective on the Primate Motor System (Oxford University Press, 2009) https://doi.org/10.1093/acprof:oso/9780195326703.001.0001.

95. X. S. J. Gershman, The Successor Representation : Its Computational Logic and Neural Substrates. 38, 7193–7200 (2018).

96. R. J. Dolan, P. Dayan, Goals and habits in the brain. Neuron 80, 312–325 (2013).

97. R. S. Sutton, A. G. Barto, Reinforcement Learning : An Introduction (MIT Press, 1998).

98. A. Berthoz, Le sens du mouvement (1997).

99. C. L. Colby, Action-oriented spatial reference frames in cortex. Neuron 20, 15–24 (1998).

100. D. M. Wolpert, S. J. Goodbody, M. Husain, Maintaining internal representations : the role of the human superior parietal lobe. Nature Neuroscience 1, 529–533 (1998).

101. W. P. Medendorp, T. Heed, State estimation in posterior parietal cortex: Distinct poles of environmental and bodily states. Progress in Neurobiology, 101691 (2019).

102. H. C. Dijkerman, W. P. Medendorp, “Visuotactile predictive mechanisms of peripersonal space” in Oxford University Press, (Oxford University Press, Oxford, 2021).

103. J. C. Cléry, S. Ben Hamed, Frontier of self and impact prediction. Frontiers in Psychology 9, 1073 (2018).

104. Z. Straka, J.-P. Noel, M. Hoffmann, A normative model of peripersonal space encoding as performing impact prediction. PLOS Computational Biology in press(2022).

105. D. H. P. Nguyen, M. Hoffmann, A. Roncone, U. Pattacini, G. Metta, Compact Real-time avoidance on a Humanoid Robot for Human-robot Interaction. 416–424 (2018).

106. J.-P. Noel, et al., Peri-personal space as a prior in coupling visual and proprioceptive signals. Scientific Reports 2018 8:1 8, 15819 (2018).

107. G. Pugach, A. Pitti, O. Tolochko, P. Gaussier, Brain-inspired coding of robot body schema through visuo-motor integration of touched events. Frontiers in Neurorobotics 13(2019).

108. D. M. Wolpert, J. R. Flanagan, Motor prediction. Current Biology 11, R729–R732 (2001).

109. W. R. Ashby, Design for a Brain: an Introduction To Cybernetics (1952).

110. K. Friston, The free-energy principle : a rough guide to the brain ? 293–301 (2009).

111. D. Lee, H. Seo, M. W. Jung, Neural basis of reinforcement learning and decision making. Annual Review of Neuroscience 35, 287–308 (2012).

112. P. Dayan, N. D. Daw, Decision theory, reinforcement learning, and the brain. Cognitive, Affective and Behavioral Neuroscience 8, 429–453 (2008).

113. A. Serino, et al., Body part-centered and full body-centered peripersonal space representations. Sci Rep 5, 18603 (2015).

114. A. Serino, E. Canzoneri, A. Avenanti, Fronto-parietal Areas Necessary for a Multisensory Representation of Peripersonal Space in Humans : An rTMS Study. 2956–2967 (2011).

115. C. L. Colby, J. R. Duhamel, Spatial representations for action in parietal cortex in *Cognitive Brain Research*, (Elsevier, 1996), pp. 105–115.

116. R. J. Bufacchi, G. D. Iannetti, The Value of Actions, in Time and Space. Trends in Cognitive Sciences 23, 270–271 (2019).

117. J.-P. Noel, A. Serino, High action values occur near the body. Trends in Cognitive Sciences (2019).

118. V. Mnih, et al., Human-level control through deep reinforcement learning. Nature 518, 529–533 (2016).

119. R. A. Andersen, V. B. Mountcastle, The influence of the angle of gaze upon the excitability of the light-sensitive neurons of the posterior parietal cortex. Journal of Neuroscience 3, 532–548 (1983).

120. R. A. Andersen, R. M. Siegel, Encoding of Spatial Location by Posterior Parietal Neurons. Science (1979) 230(1985).

121. P. R. Brotchie, R. A. Andersen, S. Lawrence H, S. J. Goodman, Head position signals used by parietal neurons to encode locations of visual stimuli. Nature 375, 232–235 (1998).

122. L. H. Snyder, K. L. Grieve, P. Brotchie, R. A. Andersen, Separate body- and world-referenced representations of visual space in parietal cortex. Nature 394, 887–890 (1998).

123. R. A. Andersen, Visual and eye movement functions of the posterior parietal cortex. Annual Review of Neuroscience 12, 377–403 (1989).

124. C. Spence, F. Pavani, J. Driver, Spatial constraints on visual-tactile cross-modal distractor congruency effects. Cogn Affect Behav Neurosci 4, 148–169 (2004).

125. C. F. Sambo, M. Liang, G. Cruccu, G. D. Iannetti, Defensive peripersonal space: the blink reflex evoked by hand stimulation is increased when the hand is near the face. J Neurophysiol 107, 880–889 (2012).

126. M. Gentilucci, C. Scandolara, I. N. Pigarev, G. Rizzolatti, Visual responses in the postarcuate cortex (area 6) of the monkey that are independent of eye position. Experimental Brain Research 50, 464–468 (1983).

127. G. Buzsáki, D. Tingley, Space and Time : The Hippocampus as a Sequence Generator. Trends in Cognitive Sciences 22, 853–869 (2018).

128. A. Barreto, et al., Transfer in Deep Reinforcement Learning Using Successor Features and Generalised Policy Improvement. Proceedings of Machine Learning Research 80(2018).

129. C. F. Sambo, G. D. Iannetti, Better safe than sorry? The safety margin surrounding the body is increased by anxiety. J Neurosci 33, 14225–14230 (2013).

130. M. Taffou, I. Viaud-Delmon, Cynophobic fear adaptively extends peri-personal space. Frontiers in Psychiatry 5, 3–9 (2014).

131. P. D. H. Nguyen, M. Hoffmann, U. Pattacini, G. Metta, “Learning peripersonal space in a humanoid robot and its application to safe human-robot interaction” (2018).

132. L. Toussaint, Y. Wamain, C. Bidet-Ildei, Y. Coello, Short-term upper-limb immobilization alters peripersonal space representation. Psychological Research 0, 0 (2018).

133. M. Bassolino, A. Finisguerra, E. Canzoneri, A. Serino, T. Pozzo, Dissociating effect of upper limb non-use and overuse on space and body representations. Neuropsychologia 70, 385–392 (2014).

134. N. X. Leclere, F. R. Sarlegna, Y. Coello, C. Bourdin, Sensori-motor adaptation to novel limb dynamics influences the representation of peripersonal space. Neuropsychologia (2019) https://doi.org/10.1016/j.neuropsychologia.2019.05.005.

